# Multivalent, Bispecific *α*B7-H3-*α*CD3 Chemically Self-Assembled Nanorings Direct Potent T-cell Responses Against Medulloblastoma

**DOI:** 10.1101/2022.03.15.484517

**Authors:** Ellie A. Mews, Pauline J. Beckmann, Mahathi Patchava, Yiao Wang, David A. Largaespada, Carston R. Wagner

## Abstract

Few therapeutic options have been made available for treating central nervous system tumors, especially upon recurrence. Recurrent medulloblastoma is uniformly lethal with no approved therapies. Recent preclinical studies have shown promising results for eradicating various solid tumors by targeting the overexpressed immune checkpoint molecule, B7-H3. However, due to several therapy-related toxicities and reports of tumor escape, the full potential of targeting this pan-cancer antigen has yet to be realized. Here, we designed and characterized bispecific chemically self-assembling nanorings (CSANs) that target the T cell receptor, CD3ε, and tumor associated antigen, B7-H3, derived from the humanized 8H9 single chain variable fragment (scFv). We show that the *α*B7-H3-*α*CD3 CSANs increase T cell infiltration and facilitate selective cytotoxicity of B7-H3^+^ medulloblastoma spheroids and that activity is independent of target cell MHC class I expression. Importantly, non-specific T cell activation against the ONS 2303 medulloblastoma cell line can be reduced by tuning the valency of the *α*CD3 targeted monomer in the oligomerized CSAN. Intraperitoneal injections of *α*B7-H3-*α*CD3 bispecific CSANs were found to effectively cross the blood-tumor barrier into the brain and elicit significant anti-tumor T cell activity intracranially as well as systemically in an orthotopic medulloblastoma model. Moreover, following treatment with *α*B7-H3-*α*CD3 CSANs, intratumoral CD4^+^ and CD8^+^ T cells were found to primarily have a central memory phenotype that displayed significant levels of characteristic activation markers. Collectively, these results demonstrate the ability of our multi-valent, bispecific CSANs to direct potent anti-tumor T cell responses and indicate its potential utility as an alternative or complementary therapy for immune cell targeting of B7-H3^+^ brain tumors.

**Significance:** This study presents *α*B7-H3-*α*CD3 bispecific CSANs as a promising, new immunotherapeutic option for patients with established B7-H3^+^ medulloblastomas.

## Introduction

Exploiting the overexpression of tumor-associated or lineage-restricted antigens on the surface of malignant cells has allowed for great strides to be made toward personalizing selective cancer therapies, specifically adoptive cell immunotherapies for hematological malignancies (1,2). However, applying this same concept to target antigens overexpressed on solid tumors is challenging due to intratumoral heterogeneity and shared expression with normal tissues. Furthermore, development of targeted immunotherapies for central nervous system (CNS) malignancies have seen even fewer advancements, particularly for treating pediatric brain tumors (3,4). Recently, chimeric antigen receptor (CAR) T cell therapies targeting highly specific or overexpressed CNS tumor antigens (e.g. EGFRviii and IL13R*α*2) have shown favorable safety profiles, however, this expression is heterogeneous between patients and tumor recurrence of antigen-negative cancer cells remains an issue (5–7).

Identification of new targetable antigens with distinct overexpression on tumor tissues has led to the discovery of B7-H3 (CD276) as a viable option for immunotherapeutic development. B7-H3 is a transmembrane protein and, similar to other B7-family members, functions as an immune modulator (8,9). The exact role of B7-H3 is still debated, though it appears that interactions with unknown immune cell ligands determine its co-stimulatory or co-inhibitory signaling (10–12). While mRNA coding for B7-H3 is found within most cells, its extracellular expression is highly regulated and relatively sparse on normal tissues; however, many groups have characterized the homogenous overexpression of B7-H3 antigen on multiple solid tumor types and the associated tumor vasculature (9,13–15). Furthermore, B7-H3 upregulation is correlated with a diminished immune response, cancer metastasis, and poor prognosis (10,16–18). Thus, B7-H3-targeting has gained increased traction in cancer immunotherapeutic development.

Two monoclonal antibodies (mAbs), 8H9 and MGA271, have shown selective and potent antitumor activity in preclinical mouse models and have since progressed into phase 1 clinical trials as Fc-enhanced antibodies, antibody-drug conjugates, and radioimmunotherapeutics (NCT03275402, NCT02982941, NCT03729596) (19–22). Additionally, preclinical results using *α*B7-H3 CAR-T cells specifically targeting CNS tumors—including neuroblastoma, medulloblastoma, and diffuse intrinsic pontine glioma (DIPG)—showed significant antitumor activity but with limited intratumoral persistence (13,14). Nevertheless, phase 1 clinical trials with *α*B7-H3 CAR-T cells are currently on-going (NCT04897321, NCT04185038). Thus, developing new B7-H3 targeted immunotherapeutics with improved tumoral infiltration and persistence could significantly advance CNS tumor treatments.

As an alternative to current mAb, bispecific T cell engager (BiTE), and CAR-T cell therapies (23,24), we have developed multi-valent, bispecific nanorings that non-genetically facilitate immune cell-to-target cell interactions. Each monomer of our chemically self-assembling nanorings (CSANs) is composed of two linked dihydrofolate reductase enzymes (DHFR^2^) fused to a targeted protein scaffold, such as a single chain variable fragment (scFv) or fibronectin (25–27). These monomers have been shown to spontaneously assemble into octameric nanorings upon addition of the chemical dimerizer, bis-methotrexate (bis-MTX), generating multi-valent CSANs (28,29). We previously demonstrated that mixing two different targeted monomers in a 1:1 ratio generates a stochastic mixture of bispecific CSANs that can be used to direct T cell interactions, via an *α*CD3 scFv, with cancer cells (**Fig. 1A**) (25). We were also able to modulate the overall CSAN affinity, and consequently T cell activity, by changing the valency of the targeted DHFR^2^ monomers in the oligomerized CSAN (30). Previous bispecific CSANs targeting CD3 and either CD22, EpCAM, or EGFR have facilitated potent T cell mediated anti-tumor activity both *in vitro* and *in vivo* (25–27).

**Figure 1.**
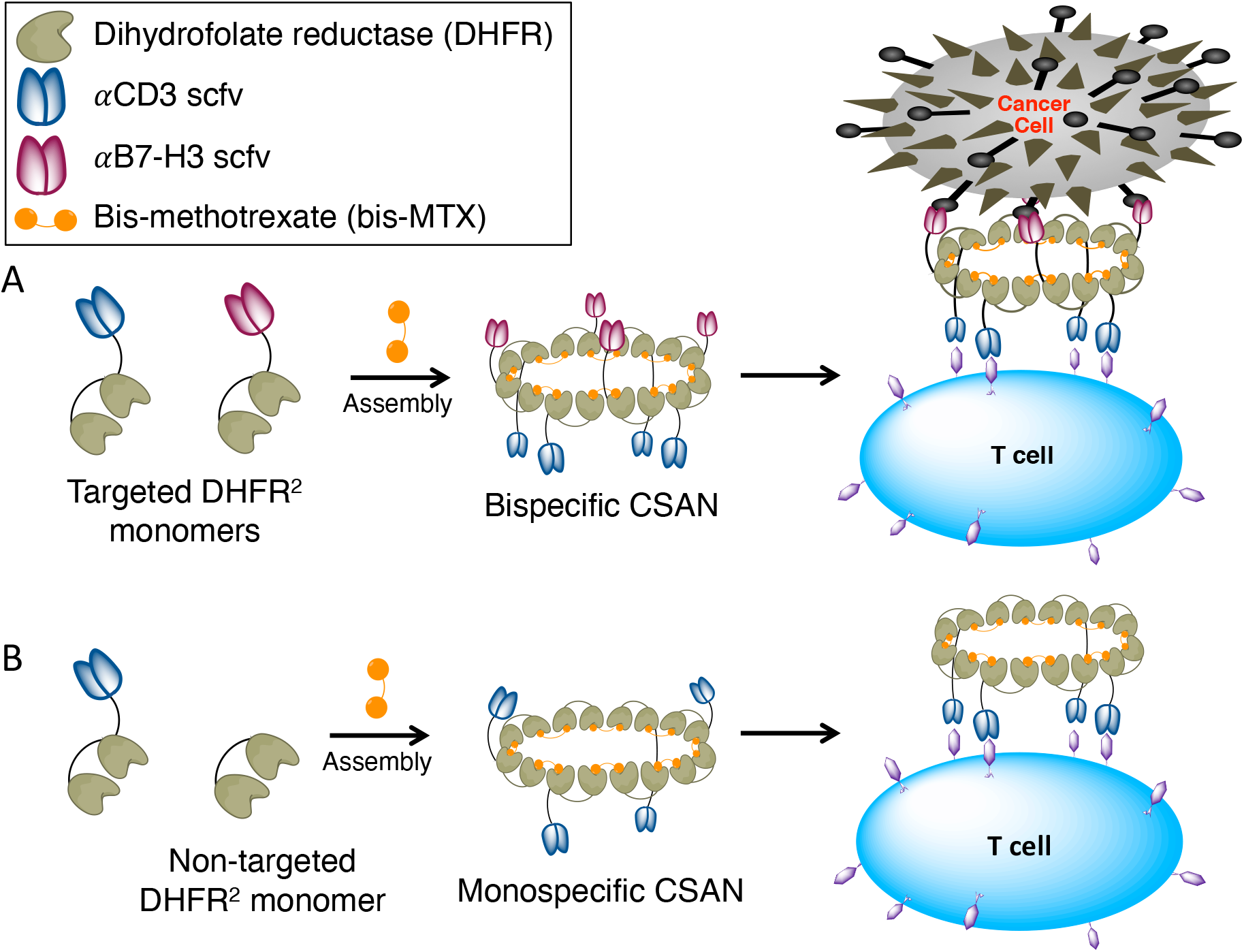
Chemically self-assembled nanorings as prosthetic antigen receptors. Bivalent DHFR^2^ protein monomers spontaneously form nanorings upon the addition of a dimeric form of methotrexate (bis-MTX). These monomers can be fused to a single *α*CD3 or *α*B7-H3 scFv, forming targeted DHFR^2^ monomers. **A**, Mixing *α*CD3 and *α*B7-H3 DHFR^2^ monomers together at a 1:1 ratio with bis-MTX forms a stochastic mixture of multivalent bispecific CSANs that can interact with both T cells and B7-H3^+^ target cells. **B**, In this work, to assess the efficacy of the bispecific CSANs, different monospecific CSANs were used as controls where only one targeted monomer—either *α*CD3 or *α*B7-H3—was mixed 1:1 with a “non-targeted” DHFR^2^ monomer to oligomerize into monospecific CSANs upon addition of bis-MTX.

To assess the potential utility of bispecific CSAN directed T cells as an immunotherapy for CNS tumors, we generated *α*B7-H3-*α*CD3 CSANs that were shown to elicit robust T cell cytotoxicity against medulloblastoma spheroids and an orthotopic xenograft model. Additionally, following intraperitoneal (IP) injections of these bispecific CSANs, we characterized the resulting T cell populations, the degree of T cell activation, and level of cytokine release both intratumorally and systemically.

## Materials and Methods

### Cell lines and culture conditions

Cell lines used in this study were provided from the following sources: human U87 MG glioblastoma by American Type Culture Collection (ATCC, Rockville, MD), human Daoy and ONS76-GFP/luciferase expressing cells by Dr. David Largaespada (University of Minnesota, Minneapolis, MN), and mouse MS1_hB7-H3_ by Dr. Ben Hackel (University of Minnesota, Minneapolis, MN). The ONS 2303 allograft cell line was created by injecting an NRG mouse cerebellum with ONS76-GFP/Luc, allowing the tumor to engraft and grow for several weeks, then removing the tumor during necropsy. The tumor was passaged through an 18G needle to create a single cell suspension and maintained in Dulbecco’s modified Eagle’s medium (DMEM) supplemented with 10% FBS, 100 U/mL penicillin, 100 μg/mL streptomycin, and 200 μg/mL hygromycin at 37°C with 5% CO_2_. The other three cancer cell lines were cultured in DMEM supplemented with 10% FBS, 100 U/mL penicillin, 100 μg/mL streptomycin at 37°C with 5% CO_2_. Human peripheral blood mononuclear cells (PBMCs) from healthy donors were isolated from the buffy coat by Ficoll density gradient centrifugation (Corning, blood obtained from Innovative Blood Resources, St. Paul, MN) and cultured in ImmunoCult-XF T cell expansion medium at 37°C with 5% CO_2_.

### Expression plasmids

A gBlock Gene Fragment coding for DHFR^2^ – *α*B7-H3 scFv fusion protein was ordered from Integrated DNA Technologies (IDT) and cloned into the Novagen pET28a(+) vector (EMD Millipore, Cat: 69864–3) via the NcoI and XhoI restriction sites. The sequence for the *α*B7-H3 scFv is a humanized version of the 8H9 scFv that was affinity matured by yeast surface display (31). Of note, the DHFR^2^ – *α*B7-H3 scFv construct has a myc tag and Flag tag to enable detection by flow cytometry for various applications.

### Protein expression and purification

The development and preparation of the DHFR^2^ – *α*CD3 scFv fusion protein monomer was previously described (29). The DHFR^2^ – *α*B7-H3 fusion protein plasmid was transformed and expressed in T7 express E. *coli* (New England Biolabs). The E. *coli* were cultured in lysogeny broth (LB) media containing 50 μg/mL of kanamycin at 37°C until the OD_600_ was within 0.6-0.8 and subsequently induced with 1mM isopropyl β-D-1-thiogalactopyranoside (IPTG) for 3 hours at 37°C. The DHFR^2^ fusion protein monomer lacking an scFv was purified from the soluble fraction of the cell lysate by IMAC as previously described (32). The scFv fusion proteins were purified from inclusion bodies as previously described (26). The scFv fusion proteins were further purified by prep-grade size exclusion chromatography (SEC) on an AKTA go (Cytiva) FPLC system. All purified proteins were analyzed by analytical size exclusion chromatography (SEC) (Cytiva) and compared to commercial molecular weight standards (Sigma Aldrich).

### CSAN assembly and characterization

CSANs were formed by adding 2 equivalents of the chemical dimerizer, bis-MTX, to a solution of DHFR^2^ fusion protein monomers in PBS (1:1:2 ratio unless specified otherwise for tuning CSAN valency). The solution was allowed to incubate for at least 30 minutes in the dark at room temperature before being characterized by analytical SEC to observe the change in hydrodynamic radius compared to the monomeric proteins. The monospecific and bispecific CSAN samples were prepared at 1 μM concentrations in PBS buffer for cryo-transmission electron microscopy (TEM) analysis as previously described (33). Images were analyzed in ImageJ and data is given as mean ± standard deviation (SD) for the indicated number of rings included in the analysis. Dynamic light scattering (DLS) and polydispersity measurements were performed on an Anton Paar Litesizer 500 (Ashland, VA), where 1mL samples (8 μM) were prepared in PBS and measured at room temperature. Hydrodynamic diameter and polydispersity values are shown as mean ± SD of three biological replicates each containing technical triplicates.

### Apparent affinity and bispecificity studies

The apparent affinity of the *α*B7-H3 monomer and *α*B7-H3 monospecific CSANs was determined by flow cytometry as previously described (32). Briefly, B7-H3 expressing cells (MS1_hB7-H3_) were incubated with different concentrations of *α*B7-H3 monomer or CSANs for 2 hours at 4°C. Cells were then washed and labeled with anti-Flag-PE antibody (Biolegend, clone L5) for 30 minutes in the dark at 4°C. The cells were then washed two times with cold PBS and analyzed on an LSR II flow cytometer (BD Biosciences). Data are displayed as the mean ± SEM of four biological replicates for each construct. The binding capability and bispecificity of the *α*B7-H3-*α*CD3 CSANs was confirmed by flow cytometry. First, the *α*CD3 scFv monomer was labeled with EZ-Link Sulfo-NHS-LC-Biotin (Thermo Fisher, 21335) according to the manufacturer’s protocol and excess NHS-biotin was removed with 0.5 mL Zeba Spin Desalting Columns (Thermo Fisher, 87766). Bispecific CSANs were then formed using the biotinylated *α*CD3 scFv monomer and the *α*B7-H3 scFv monomer at a 1:1 ratio. MS1_hB7-H3_ was selected as the B7-H3^+^/CD3^−^ cell line and human PBMCs were selected as the B7-H3^−^/CD3^+^ cell line. The cells were incubated with 100nM bispecific CSANs for 1 hour at 4°C, after which the cells were washed and labeled with different fluorescent secondaries for 30 minutes in the dark at 4°C. To detect the presence of the *α*B7-H3 portion of the ring on the PBMCs, the cells were labeled with anti-Flag-PE antibody. To detect the presence of the *α*CD3 portion of the ring on MS1_hB7-H3_, the cells were labeled with streptavidin-Alexa Fluor 647 (AF647). The cells were subsequently washed with cold PBS and analyzed using an LSR II flow cytometer. The mean fluorescence intensity (MFI) was monitored and compared to unstained and control samples.

### Quantification of B7-H3 and MHC class II antigen expression

The cells used in this study were labeled with anti-human B7-H3 monoclonal antibody (Biolegend, clone MIH42) PerCP/Cyanine 5.5 conjugate at 4°C for 45 minutes in the dark. Cells were then washed twice and analyzed on a BD LSR II cytometer. This MFI value was compared to a calibration curve prepared from Quantum Simply Cellular anti-mouse IgG calibration beads (Bangs Laboratories, 815) labeled with the same B7-H3 conjugated antibody according to the manufacturer’s protocol. Values are displayed as the mean ± SD from three technical replicates. Similarly, ONS 2303 cells and the IgG calibration beads were stained with anti-human HLA-DR, DP, DQ monoclonal antibody (Biolegend, clone Tü39) at 4°C for 45 minutes in the dark. Cells and beads were then washed and stained with goat anti-mouse secondary antibody (Cat: A-21235, AF647) at 4°C for 30 minutes in the dark. Cells and beads were then washed twice and analyzed on a BD LSR II cytometer.

### T cell binding and infiltration microscopy

To observe T cells binding to a monolayer of target cells, 1×10^5^ ONS 2303 cells and 1.8×10^5^ MS1_hB7-H3_ cells were allowed to adhere to glass coverslips overnight in 6 well plates. After adhering, the MS1 cells were labeled with CFSE (Biolegend, 423801) according to manufacturer’s protocol and then both cell types were washed and the coverslips were blocked with PBS + 5% BSA solution for 45 minutes at room temperature. During this time, rested PBMCs were labeled with Cell Trace Far Red Dye (Thermo Fisher, C34564) according to manufacturer’s protocol and then washed. After blocking, the target cells were washed and then pre-incubated with 300 nM CSANs or PBS for 30 minutes at 37°C. After which the labeled PBMCs were added to each well at a 10:1 effector to target ratio (E:T), without washing, and the co-culture was incubated for 2 hours at 37°C. The coverslips were washed 3 times by PBS with gentle nutation for 5 minutes to remove any unbound T cells and CSANs. The bound cells remaining were fixed in 4% paraformaldehyde for 15 minutes at room temperature and then washed twice with PBS. The coverslips were removed from the plates, rinsed with ultrapure water, and mounted on glass slides with Vectashield antifade mountant without dapi (Vector Leboratories) for 24 hours at room temperature in the dark. All images were taken on a Nikon A1R FLIM Confocal Microscope (Nikon Instruments) using a 20x water immersion objective. Confocal microscopy performed on ONS 2303 spheroids was performed similarly. 0.2×10^5^ ONS 2303 cells were plated in 96 well ultra-low attachment (ULA) U-bottom plates and centrifuged; spheroids formed within 2 days. Rested PBMCs were labeled with far red dye and the control and treatment CSANs were oligomerized as described above. The media in the ULA plate was replaced with fresh DMEM and then spheroids were pre-incubated with 100 nM CSANs or PBS for 30 minutes at 37°C. After which the labeled PBMCs were added to each well at a 20:1 E:T ratio, without washing, and the co-culture was incubated for either 30 minutes or 2 hours at 4°C. After incubation, the contents of the entire wells were moved into 1.5 mL eppendorf tubes and washed with cold PBS three times. The spheroids and bound T cells were fixed with 4% paraformaldehyde for 15 minutes at room temperature, washed with PBS twice, washed with ultrapure water once, and then cleared using CytoVista 3D Cell Culture Clearing Reagent (Thermo Fisher, V11326) according to manufacturer’s protocol. The clearing reagent was carefully removed and the spheroids were mounted with Vectashield into a chambered μ-Slide (Ibidi, 81506) for 24 hours at room temperature in the dark. All images were taken on a Nikon A1R FLIM Confocal Microscope (Nikon Instruments) using a 20x water immersion objective. Spheroids were penetrated at least 100 μm to create z-projection images. All microscopy images were analyzed with ImageJ software and data is shown as mean ± SEM with sample sizes indicated in the figure legend.

### Cytotoxicity assays

Real time visualization of cancer cell death was monitored by loss of ONS 2303 GFP fluorescence in an Incucyte SX5. One day prior to the assay, 5 × 10^3^ ONS 2303 target cells were seeded into a 96 well plate in 100 μL of complete DMEM media per well. The human PBMCs were thawed and allowed to rest in media overnight. 16 hours later, media in the plate was replaced with fresh DMEM, the unactivated PBMCs were counted, and monospecific and bispecific CSANs treatments were oligomerized. CSANs or PBS (50 μL per well of the indicated concentration) and the appropriate number of PBMCs (50 μL per well) were added to the plate at the same time (this number is determined by the indicated E:T ratio). The plate was placed in the Incucyte SX5 at 37°C and 5% CO_2_ and GFP fluorescence was monitored for 72 hours (the PBMCs were not fluorescent or stained). The ONS 2303 count was calculated using the SX5 Cell-by-Cell module. Cytotoxicity assays performed on ONS 2303 spheroids were performed similarly. Just before placing spheroids in the Incucyte, 2.5% v/v cold Matrigel was added to the top of each well. The total spheroid area was calculated using the SX5 spheroid module. All Incucyte images were obtained with a 10x objective and each cytotoxicity graph (n=3 wells per time point) was obtained using one PBMC donor but is representative of three different donors and is shown as mean ± SEM calculated by the Incucyte software. Normalized data is shown as a ratio of the ONS 2303 viability compared to viability at time zero also calculated by the Incucyte software.

### Cytokine assays

The supernatants from effector and target cell co-cultures were collected from separate 24 and 72 hour plates. IL-2 and IFN*γ* cytokine concentrations were analyzed by sandwich ELISA according to manufacturer’s kit protocol (BD Opteia human IL-2 and human IFN*γ*). *In vivo* cerebellum lysates were analyzed for cytokines similarly. Intracranial samples were collected from inoculated mouse cerebellum as described below. The flash frozen piece of cerebellum was lysed with 300 μL lysis buffer (RayBiotech, EL-lysis), sonicated for 2 seconds, and centrifuged to remove any remaining solids. Total protein in the lysates were analyzed by BCA assay (Genesee Scientific, 18-440) and normalized before use in the ELISA assays as described.

### Antibodies and flow cytometry studies

Staining for CSANs that remained bound to effector/target cells 24 or 72 hours after co-culture began was performed with anti-Flag-PE antibody. PBMCs and target cells from the 24 or 72 hour co-cultures were determined by forward/side scatter and dead cells were removed using Ghost Dye Red 780 (Tonbo biosciences). T cell activation markers were stained with anti-human CD4 (BD Biosciences, clone SK3, BUV395), anti-human CD8 (Biolegend, clone SK1, BV711), anti-human CD25 (Biolegend, clone BC96, PE/Dazzle 594), and anti-human CD107a (Biolegend, clone H4A3, PE). Monensin (BD Biosciences) was added 2-4 hours prior to staining. Memory T cell analysis was performed using anti-human CD4, anti-human CD8, anti-human CCR7 (Biolegend, clone G043H7, AF647), and anti-human CD45RO (Biolegend, clone UCHL1, BV421) on the co-cultured cells 24 or 72 hours after the assay began. All flow cytometry prep was done at 4°C in FACS buffer (PBS + 0.1% BSA), analysis was done using an LSRFortessa H0081 (BD Biosciences), and values are displayed as the mean ± SEM of three biological replicates each containing technical triplicates. Intracranial samples were collected from inoculated mouse cerebellum as described below. The piece of cerebellum was dissociated into a single cell suspension by a gentleMACS Octo Dissociator within a gentleMACS C Tube in combination with the Brain Tumor Dissociation Kit (P) (Miltenyi Biotec) following all associated manufacturer’s protocols. Systemic blood samples were also collected from these mice as described below. Analysis of *in vivo* samples was then performed similarly to that described above for CSAN detection, T cell activation marker, and memory T cell analyses. ONS 2303 GFP expression allowed for the quantitation of remaining cancer cells in the intracranial samples. Intracranial regulatory T cell response was analyzed using anti-human CD4, anti-human CD25, and anti-human Foxp3 (Biolegend, clone 206D, AF488) antibodies. The cells were surface stained with CD4 and CD25, then fixed and permeabilized using eBioscience Foxp3/Transcription Factor Staining Buffer Set (Thermo Fisher), and then stained for Foxp3. Finally, intracranial and systemic lymphocyte samples were stained with anti-human CTLA-4 antibody (Biolegend, clone BNI3, BV421). Additionally, Precision Count Beads (Biolegend) were added to the *in vivo* samples to obtain absolute counts of events acquired on the cytometer as well as using human and mouse Fc blocking antibodies (Biolegend) to reduce non-specific antibody staining. All flow cytometry prep was done at 4°C in FACS buffer and analysis was done using an LSRFortessa H0081. Values are displayed as the mean ± SEM of 5-7 mouse samples for each treatment type.

### Orthotopic medulloblastoma tumor model

NRG mice (NOD.Cg-*Rag1*^*tm1Mom*^ *Il2rg*^*tm1Wjl*^/SzJ, Jackson 007799) were injected as previously described (34). Briefly, ONS 2303-GFP/Luc cells were prepared in PBS and stored on ice prior to injection. Mice were anesthetized (54 mg/mL ketamine, 9.2 mg/mL xylazine) and injected with 1×10^5^ ONS 2303 cells in 5 μL over 5 minutes (stereotactic coordinates: 2.5mm posterior to lambda, 1.5mm to the right, with needle angled 2.5° to the right). Successful injection and engraftment was verified by luciferase imaging using the Xenogen IVIS-100 Imaging System (PerkinElmer) as previously described (35). 15 days after tumor inoculation, 20×10^6^ unactivated human PBMCs were intravenously (IV) injected. 4 days after PBMCs were given, mice received intraperitoneal (IP) injections of 1mg/kg CSANs or PBS. Booster injections of CSANs or PBS were given every other day for 5 total treatments; the 5^th^ treatment was staggered for some mice so that the flow cytometry analysis on live cells would happen about 24 hours after the final injection. IVIS imaging and body weight was monitored twice per week. Mice were sacrificed at about day 30 and the blood and cerebellum were collected for analysis. The blood was placed on heparin (BD Biosciences, 365965), red blood cells were removed using ACK lysing buffer, and the remaining lymphocytes were washed and used for flow cytometry analysis. The cerebellum was sliced in half: one half was flash frozen and stored at −80°C until cytokine analysis as described above, the other half was dissociated into a single cell suspension and used in flow cytometry analysis as described above. IVIS and body weight data is shown as mean ± SEM of 5-7 mouse samples for each treatment type.

### CRISPR knock-down of MHC class I

Knock down (KD) of MHC class I expression was achieved by targeting β_2_ microglobulin (*B2M*, P61769) using sgRNAs selected in the Synthego Knockout Guide Design Tool. ONS 2303 cells were cultured to 70% confluency, harvested, and pelleted. Diluted sgRNAs were mixed with Cas9 at a 3:1 ratio and incubated at room temperature for 10 minutes to form RNPs. The RNPs were then resuspended in R buffer (Invitrogen, MPK1025) and transferred to pelleted cells. The cell and RNP mixture was electroporated using a Neon Transfection System (Invitrogen) according to manufacturer’s protocol. The electroporated cells were immediately placed in a pre-warmed 12 well plate with complete DMEM. The cells were expanded for one week and those that were double negative for MHC class I (Biolegend, W6/32, AF700) and β_2_ microglobulin (Biolegend, clone 2M2, PE) were selected via fluorescence assisted cell sorting (FACS). Collected cells were passaged no more than 3 times prior to cytotoxicity analysis and the expression of MHC class I was checked on the same day as shown (Supplementary Fig. S6). Wild type (WT) and KD cells were plated for cytotoxicity assays as described above. The plate was placed in the Incucyte Zoom with 10x objective at 37°C and 5% CO_2_ and GFP fluorescence was monitored for 72 hours (the PBMCs were not fluorescent or stained). Data is shown as mean ± SEM (n=3 wells per time point) calculated by the Incucyte software.

### Statistical Analysis

Data was analyzed using two-tailed unpaired Student’s t-test for comparison of treatment to control groups. P < 0.05 was considered statistically significant and values are denoted with asterisks as follows: ns P > 0.05 not significant; * P < 0.05; ** P < 0.01; *** P < 0.001 or value is indicated within the figure and legend.

## Results

### Preparation and characterization of αB7-H3 DHFR^2^ fusion protein

We have previously reported that our bispecific CSANs are able to direct T cell cytotoxicity against solid tumor targets both *in vitro* and *in vivo* (26,27). To expand the therapeutic capacity of our nanorings, we investigated their ability to cross the blood-tumor barrier (BTB) and mediate a T cell response against established brain tumors. B7-H3 has been shown to be broadly overexpressed in many different cancer types, particularly pediatric medulloblastoma, but has limited expression on normal tissues (13–15). Therefore, we developed *α*B7-H3-*α*CD3 bispecific CSANs to characterize their impact on T cell infiltration, activation, and cytotoxicity *in vitro* and *in vivo* against an allograft of ONS76 medulloblastoma cells, which is referred to as ONS 2303.

Similar to the design of the *α*CD3 DHFR^2^ monomer that was previously reported, a humanized *α*B7-H3 scFv originating from the monoclonal antibody, 8H9, was fused to the C-terminus of the DHFR^2^ monomer and expressed in E. *coli* as an insoluble protein (29,31). As controls, we also prepared two separate *α*B7-H3 and *α*CD3 monospecific CSANs in which the corresponding targeted monomer is replaced with a “non-targeted” DHFR^2^ monomer (**Fig. 1B**). Following the expression of both targeted monomers, size exclusion chromatography (SEC) was used to confirm their purity and characterize the formation of *α*B7-H3-*α*CD3 bispecific CSANs after incubation with the chemical dimerizer, bis-MTX (**Fig. 2A**). As observed for previously reported bispecific CSANs, the oligomerized *α*B7-H3-*α*CD3 nanorings eluted faster due to their larger hydrodynamic radius compared to the smaller monomers (26,27). Additionally, dynamic light scattering (DLS) showed that upon oligomerization, *α*B7-H3 monospecific nanorings had a larger hydrodynamic diameter (15.8 ± 0.7 nm) compared to the monomers themselves (*α*B7-H3 monomer at 9.4 ± 0.2 nm, *α*CD3 monomer at 9.2 ± 0.3 nm) (**Fig. 2B and Supplementary Fig. S1A**), which was also observed in previous CSAN reports (26,30). Importantly, the oligomerized CSANs fall within an acceptable polydispersity index range at less than 25%, which is considered to be a uniform sample with respect to particle size and is important for delivery across the blood-brain barrier (BBB) (**Supplementary Fig. S1B**) (36). The CSAN structure was further confirmed by directly visualizing the nanorings using cryo-transmission electron microscopy (TEM). The average diameter of the *α*B7-H3-*α*CD3 bispecific nanorings was found to be 22 ± 3 nm (**Fig. 2C**) and the diameter of the *α*B7-H3 monospecific nanorings was 20 ± 3 nm (**Supplementary Fig. S1C**), which is consistent with previous CSAN characterization studies (26,37). As predicted, the affinity titration experiments demonstrated that upon oligomerization the apparent affinity of the monomer (139 ± 14 nM) increases 4.5-fold as the valency of the targeting elements incorporated in the CSAN increases (31 ± 1 nM) (**Fig. 2D**).

**Figure 2.**
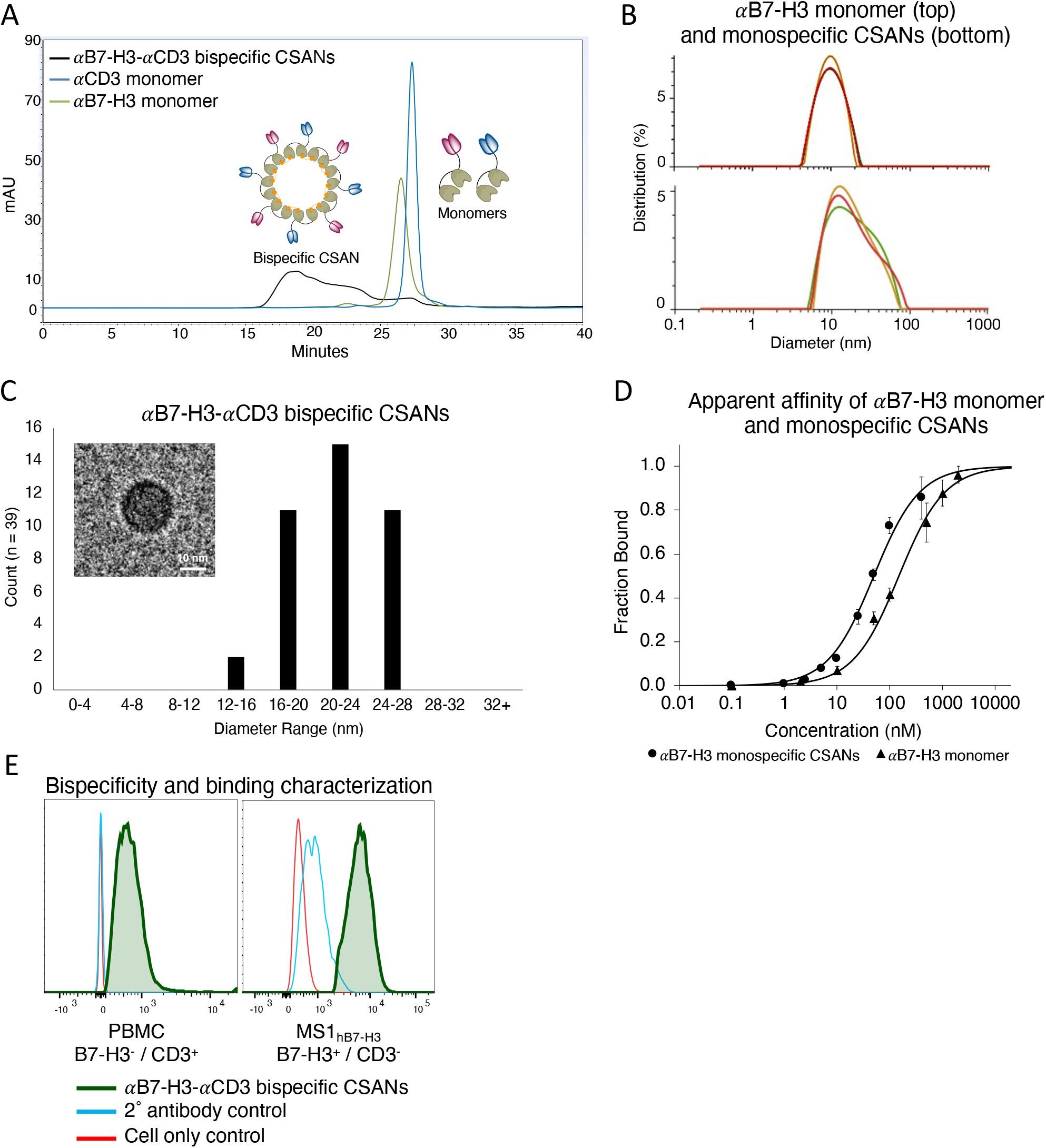
*α*B7-H3 CSAN characterization. **A**, Size exclusion chromatography confirmed monomeric *α*B7-H3 and *α*CD3 purity as well as *α*B7-H3-*α*CD3 bispecific CSAN formation upon addition of bis-MTX. **B**, The hydrodynamic diameter of the *α*B7-H3 monomer, top (9.4 ± 0.2 nm), and *α*B7-H3 monospecific CSANs with an average of 4 targeting elements, bottom (15.8 ± 0.7 nm), was determined using DLS. Representative images of 3 experiments shown. **C**, Cryo-TEM size distribution analysis (average = 22 ± 3 nm) and a representative image of *α*B7-H3-*α*CD3 bispecific CSANs. Scale bar, 10nm. **D**, Flow cytometry was used to determine the apparent affinity of *α*B7-H3 monomer (139 ± 14 nM) and *α*B7-H3 monospecific CSANs with an average of 4 targeting elements (31 ± 1 nM) using human B7-H3 stably transfected murine endothelial cells, MS1_hB7-H3_. (Data is displayed as mean ± SEM, n = 3 biological replicates). **E**, The bispecificity and binding capability of the *α*B7-H3-*α*CD3 bispecific CSANs was confirmed by flow cytometry. Using PBMCs as the CD3^+^ only target, a shift in fluorescence to the right indicates the presence of B7-H3 targeted monomer in the bispecific CSAN; and using MS1_hB7-H3_ as the B7-H3^+^ only target, a shift in fluorescence to the right indicates the presence of CD3 targeted monomer in the bispecific CSAN. Because both targeting elements are detected on both cell types, the nanorings are confirmed to be bispecific and functional (green trace).

Upon confirmation of ring formation, the bispecificity and binding functionality of the *α*B7-H3-*α*CD3 CSANs was tested against a B7-H3^+^ cell line, MS1_hB7-H3_, and CD3^+^ peripheral blood mononuclear cells (PBMCs). The *α*B7-H3 monomer was expressed with a FLAG tag to enable fluorescent detection by flow cytometry via an *α*FLAG antibody, whereas the *α*CD3 monomer was not designed with a protein tag and was instead non-specifically labeled with NHS-biotin to enable detection via fluorescently conjugated streptavidin. After removal of unreacted NHS-biotin, the two monomers were oligomerized into bispecific and corresponding monospecific CSANs and incubated with the B7-H3^+^ MS1_hB7-H3_ cells or CD3^+^ PBMCs. Monospecific NHS-labeled *α*CD3 CSANs were not detected on the surface of MS1_hB7-H3_ cells because they do not express CD3 antigen (**Supplementary Fig. S1D**). Similarly, the monospecific *α*B7-H3 CSANs were not detected after incubation with the B7-H3^−^ PBMCs. However, when incubated with the bispecific nanorings, NHS-labeled *α*CD3 is detected on the surface of MS1_hB7-H3_ cells and the FLAG-tagged *α*B7-H3 is detected on the PBMCs, indicating that the CSANs are indeed bispecific and functional (**Fig. 2E**).

### Bispecific CSANs increase T cell binding and infiltration of 2D and 3D cultures in vitro

As previously discussed, many different patient derived tumors express B7-H3, with notable overexpression in pediatric brain tumors (13–15). We sought to validate the degree this expression changes when using different culturing techniques on various immortalized cancer cell lines. We found that when cultured as a 2D monolayer, the medulloblastoma cell lines (ONS 2303 and Daoy) expressed significantly higher levels of B7-H3 antigen compared to when cultured as a 3D spheroid (**Fig. 3A**). The mouse cell line, MS1_hB7-H3_, was transduced to express greater than 10^6^ human B7-H3 antigens per cell and was used as a positive control. MS1 expression of B7-H3 antigens in 3D culture could not be made as these cells do not form spheroids under standard conditions.

**Figure 3.**
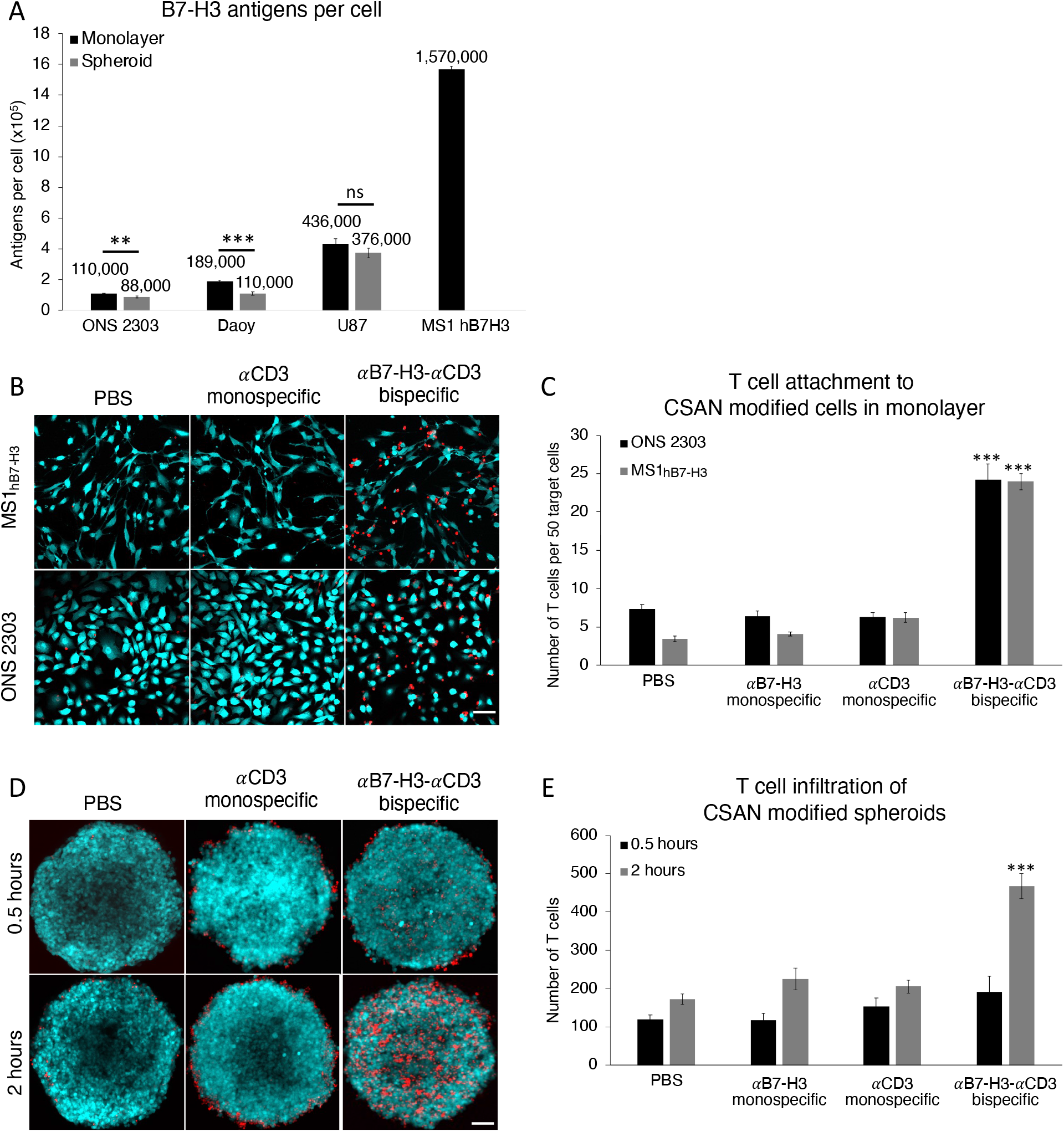
*α*B7-H3-*α*CD3 bispecific CSANs increase T cell infiltration and attachment to B7-H3^+^ cells modified with CSANs. **A**, Quantification of B7-H3 antigen on different immortalized brain cancer cells cultured using different methods and the stably transfected MS1_hB7-H3_. (Data is displayed as mean ± SD, n = 3 technical replicates). A 2D monolayer of ONS 2303 allograft or MS1_hB7-H3_ cells were pre-incubated with monospecific or bispecific CSANs for 30 minutes at 37°C, after which unactivated T cells were added and the entire co-culture was incubated for 2 hours at 37°C. **B**, Representative composite fluorescent images of the co-culture for each cell line. Scale bar, 50μm. **C**, The number of T cells that remained attached to the monolayer after the 2 hour incubation were quantified using fluorescence microscopy (Data is displayed as mean ± SEM, *** P<0.0001 with respect to PBS and monospecific control CSANs by two-tailed unpaired t-test, n = 7-12 images each). 3D spheroids of ONS 2303 allograft cells were pre incubated with monospecific or bispecific CSANs for 30 minutes at 37°C, after which unactivated T cells were added and the entire co-culture was incubated for either 0.5 or 2 hours at 4°C. **D**, Representative z-projection images of the co-culture for each incubation time. Scale bar, 50μm. **E**, Z-stack images were acquired using fluorescent confocal microscopy to quantify the T cells that attached to and infiltrated the spheroids at the different incubation times (Data is displayed as mean ± SEM, *** P<0.0001 with respect to PBS and monospecific control CSANs by two-tailed unpaired t-test, n = 5-6 spheroids each).

To assess the effect bispecific CSANs had on T cell interactions with high expressing (MS1_hB7-H3_) and low expressing (ONS 2303) target cells, we quantified the number of T cells that bound to a monolayer of target cells that had been pre-incubated with either controls or *α*B7-H3-*α*CD3 CSANs (300 nM). After removing any unbound cells and unbound CSANs, the remaining fluorescently labeled T cells were visualized by fluorescent microscopy bound to either CFSE-labeled MS1_hB7-H3_ cells or GFP-expressing ONS 2303 cells (**Fig. 3B**). The *α*B7-H3-*α*CD3 bispecific CSANs significantly increased T cell binding compared to non-targeted controls by 3-fold, regardless of the quantity of B7-H3 antigen expression (**Fig. 3C**). Similarly, the ability of the bispecific CSANs to augment T cell binding and infiltration of medulloblastoma spheroids over time was characterized by fluorescent confocal microscopy with images collected to a depth of at least 100 μm (**Fig. 3D**). Interestingly, T cell binding and infiltration of ONS 2303 spheroids was significantly improved when directed with *α*B7-H3-*α*CD3 CSANs and appeared to be time dependent (**Fig. 3E**).

### Bispecific CSANs facilitate T cell cytotoxicity in 2D and 3D cultures in vitro

*α*B7-H3-*α*CD3 CSAN directed T cell cytotoxicity was first investigated on a monolayer of ONS 2303 target cells that were co-cultured with unactivated PBMCs and treated with variable concentrations of CSANs over a 72 hour period (**Fig. 4A** and **Supplementary Fig. S2A-C**). Directed T cell cytotoxicity was dose dependent with significant target cell death observed at the lowest CSAN concentration (50nM) and maximum T cell killing observed at the highest CSAN concentration (400 nM) compared to the PBS control. Using a similar co-culture, we analyzed different effector-to-target (E:T) ratios at a fixed concentration of CSANs (400 nM) and observed significant targeted cell killing at as low as a 1:1 ratio and up to a 6-fold difference in cytotoxicity at the largest E:T ratio (20:1) using bispecific CSANs compared to the PBS control (**Fig. 4B**). Interestingly, the *α*CD3 monospecific CSANs appeared to elicit a similar level of non-targeted T cell cytotoxicity against the ONS 2303 cells.

**Figure 4.**
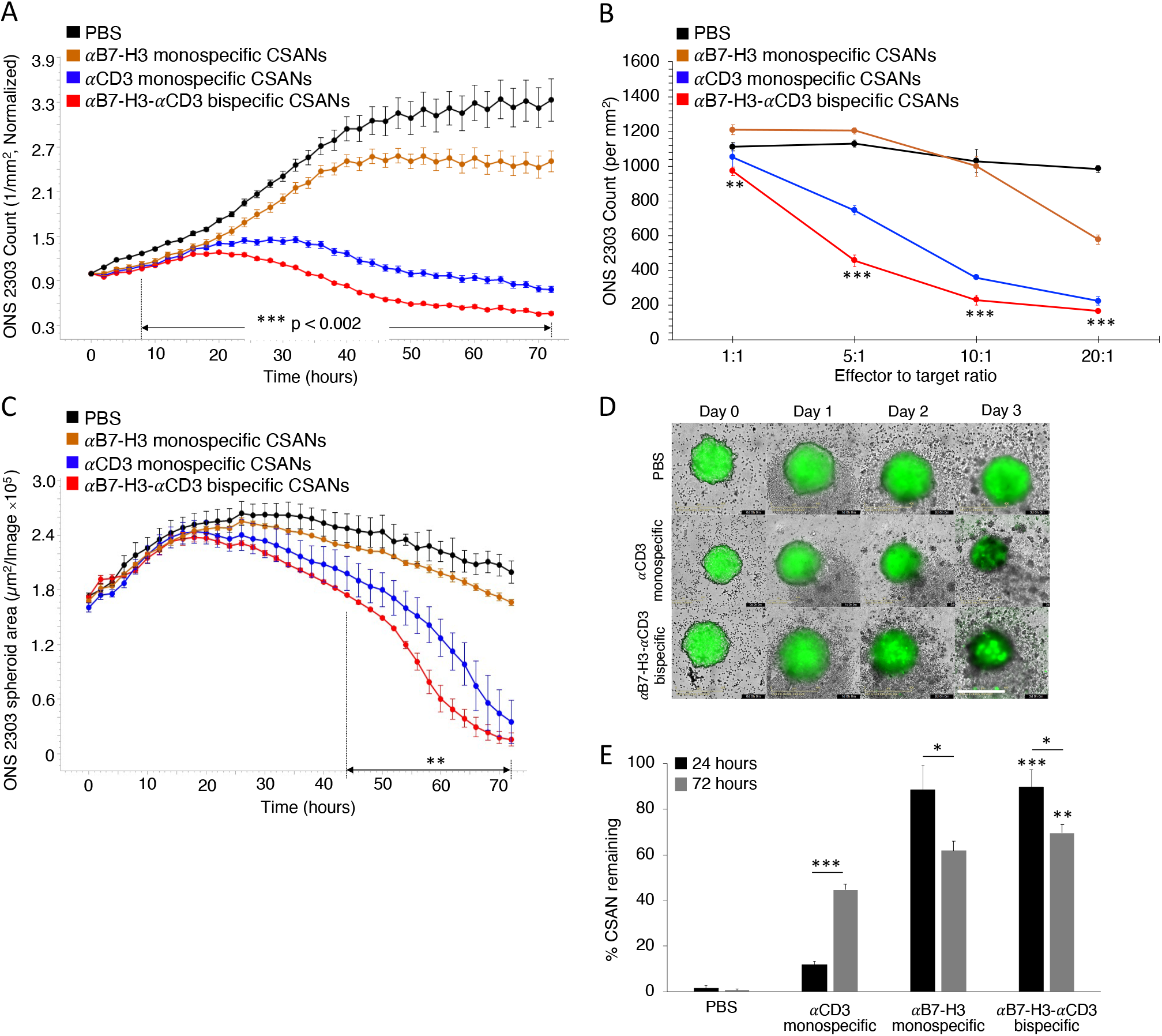
*α*B7-H3-*α*CD3 bispecific CSANs direct selective T cell cytotoxicity against B7-H3^+^ target cells. ONS 2303 was seeded into a plate and cultured as a monolayer. Unactivated PBMCs were then added to the target cells 16 hours later with either monospecific or bispecific CSANs all at once. **A**, ONS 2303 cell viability was monitored over a 72 hour period using an Incucyte SX5 at a fixed CSAN concentration (400nM) and PBMCs added at a 20:1 E:T ratio (***P<0.002 with respect to PBS by two-tailed unpaired t-test). **B**, Different E:T ratios were similarly analyzed at a fixed CSAN concentration (400nM) where each bullet point represents the final ONS 2303 viable cell count at the end of the 72 hour cytotoxicity study (**P<0.01, ***P<0.001 with respect to PBS by two-tailed unpaired t-test). **C**, ONS 2303 was seeded into a 96 well ULA plate and cultured as a spheroid for 2 days. Unactivated PBMCs were then added to the spheroids with either monospecific or bispecific CSANs all at once. The spheroid area was monitored over a 72 hour period using an Incucyte SX5 at a fixed CSAN concentration (400nM) and PBMCs added at a 5:1 E:T ratio (**P<0.01 with respect to PBS by two-tailed unpaired t-test). **D**, Representative images of ONS 2303 spheroids from **C** treated with PBMCs and either PBS, *α*CD3 monospecific, or *α*B7-H3-*α*CD3 bispecific CSANs over time. Scale bar, 400μm. **E**, A similar monolayer co-culture was incubated for 24 hours or 72 hours before removing from the plate and measuring CSAN (Flag tag) presence on target/effector cell surfaces by flow cytometry (*P<0.05, **P<0.01, ***P<0.001 with respect to PBS or *α*CD3 monospecific CSAN control for each individual data series by two-tailed unpaired t-test or as indicated by line marker). Incucyte data was obtained using one PBMC donor but is representative of three different donors (**Supplementary Fig. S3 and S4**) (n = 3 wells per time point ± SEM). All Incucyte images were taken with a 10x objective. Normalized data is shown as a ratio of the ONS 2303 viability compared to viability at time zero. For the flow cytometry experiment, effector and target cells were determined by forward/side scatter and dead cells were removed using a viability dye (n = 9 using 3 PBMC donors total ± SEM).

Because the culture technique was shown to affect the expression of B7-H3 antigens per cell, we assessed whether this change in expression also affected the cytotoxic potential of T cells directed with *α*B7-H3-*α*CD3 CSANs. Using a similar 72 hour live cell imaging assay, the ONS 2303 spheroids were treated with controls or bispecific CSANs and a 5:1 E:T ratio of unactivated PBMCs. A significant dose and time dependent reduction in spheroid size was observed for samples treated with targeted bispecific CSANs compared to the PBS control (**Fig. 4C** and **Supplementary Fig. S2D**). Similar to the 2D cytotoxicity results, the *α*CD3 monospecific CSANs directed comparable T cell killing against the ONS 2303 spheroids as that observed for targeted *α*B7-H3-*α*CD3 CSANs. Although, cytotoxicity was observable only after day two, possibly reflecting the time dependence required for spheroid infiltration by the T cells (**Fig. 4D** and **Supplementary Fig. S2E**).

To ascertain the longevity of the *α*B7-H3-*α*CD3 bispecific CSANs bound to the surfaces of target and effector cells, flow cytometry was used to detect the nanorings near the beginning of the co-culture at 24 hours and the end at 72 hours. There were significantly more bispecific CSANs detected at both time points compared to the PBS and *α*CD3 monospecific CSAN controls (**Fig. 4E**). However, the amount of *α*B7-H3 monospecific CSANs bound to the ONS 2303 cells was similar at 24 and 72 hours to that of the bispecific nanorings. This is likely due to the tight binding affinity of the *α*B7-H3 scFv which is further enhanced by the increased scFv avidity in the multivalent CSAN construct. Interestingly, the percentage of cells bound to *α*CD3 monospecific CSANs increased over time perhaps due to the necessity for increased exposure time to CD3 antigen. Notably, the detectable amount of bispecific CSANs did decrease between 24 (90%) and 72 hours (70%) but was still reasonably high indicating that bispecific nanorings stably bind B7-H3 antigen and possibly undergo minimal internalization.

### Effect of bispecific CSANs on T cell activation and memory cell formation

The cytotoxicity caused by the bispecific CSAN modified PBMCs was corroborated by the cytokine release profiles present in the 24 and 72 hour co-culture supernatants. PBMC production of IFN*γ* and IL-2 was already significant at 24 hours (1.9-fold and 2.3-fold higher compared to the *α*CD3 monospecific control, respectively) when treated with *α*B7-H3-*α*CD3 CSANs and continued to increase over the course of 72 hours (**Fig. 5A-B**). This cytokine response was shown to be dose dependent with regard to *α*B7-H3-*α*CD3 CSANs, particularly after 72 hours of co-culture (**Supplementary Fig. S2F-G**) correlating well with the bispecific CSAN directed T cell cytotoxicity discussed above. Additionally, we investigated the ability of *α*B7-H3-*α*CD3 CSANs to specifically activate CD8^+^ T cells by measuring expression of the degranulation marker, CD107a, and late-activation marker, CD25, when co-cultured with ONS 2303 target cells for 24 or 72 hours. Consistent with the cytokine release profiles, expression of CD107a and CD25 was significant at 24 hours (2.4-fold and 2.3-fold higher compared to the *α*CD3 monospecific control, respectively) and continued to increase through the 72 hour co-culture when treated with bispecific CSANs (**Fig. 5C-D**). These observations are comparable to different targeted CSANs that have been previously reported (25–27). In addition, similar levels of IFN*γ* and IL-2 secretion and CD107a expression were observed by others utilizing *α*B7-H3 CAR-T cells targeting various solid tumor types *in vitro* (13,14,38,39). These results suggest that, when modified with the *α*B7-H3-*α*CD3 bispecific CSANs, T cell activation is sustained and increases over at least a 72 hour period.

**Figure 5.**
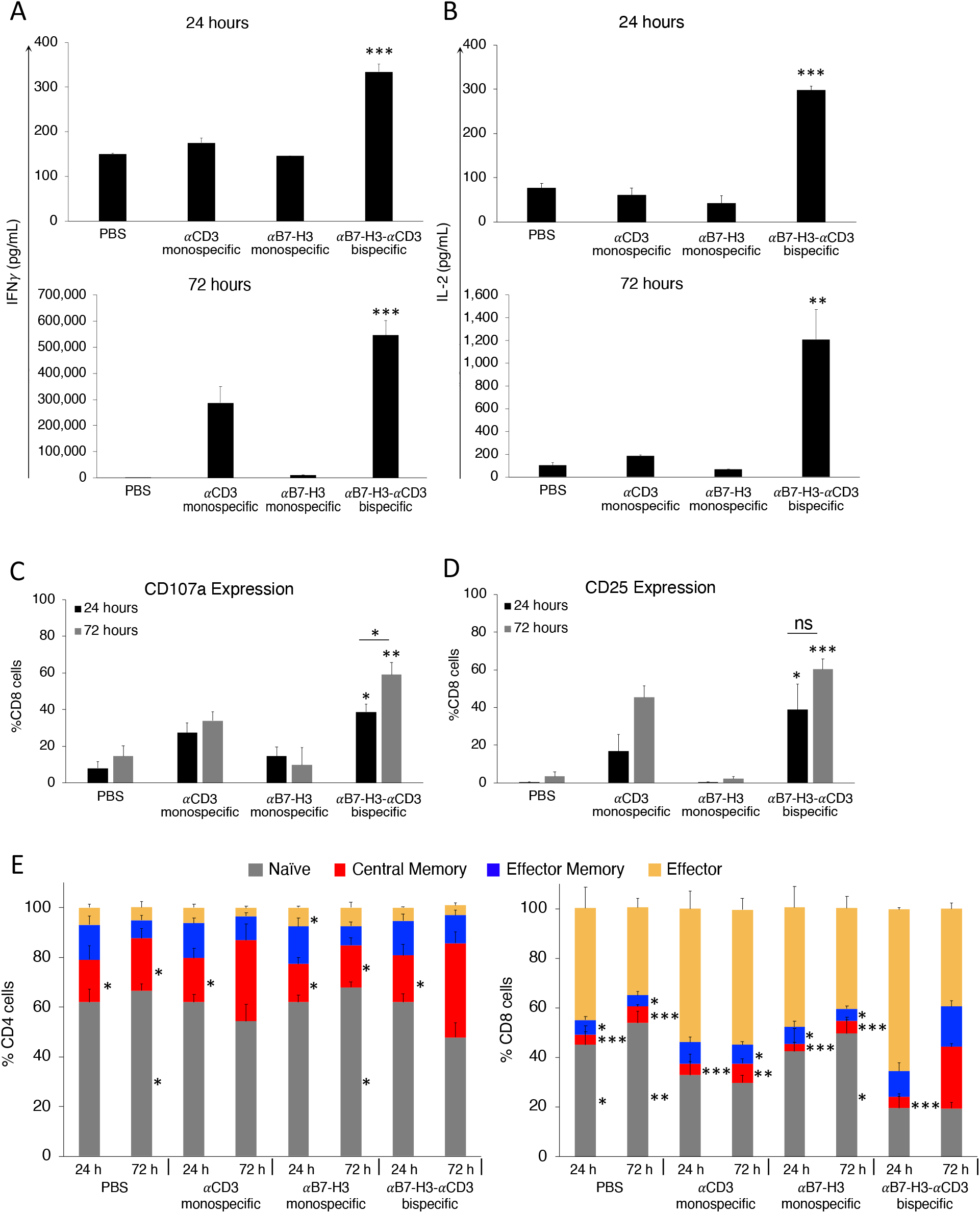
*In vitro* flow cytometry investigation of *α*B7-H3-*α*CD3 CSAN effect on T cell activation and population changes over time. Following the cytotoxicity study, at the specified time points, the media from the co-culture was analyzed for **A**, IFN*γ* and **B**, IL-2 using a sandwich ELISA (*P<0.05, ***P<0.001 with respect to PBS and *α*B7-H3 monospecific CSAN controls by two-tailed unpaired t-test). A similar co-culture was set up for either 24 or 72 hours where golgistop (monensin) was added 2-4 hours prior to cell harvest and then surface expression of **C**, CD107a and **D**, CD25 was determined on CD8^+^ T cells by flow cytometry (*P<0.05, ***P<0.001 with respect to PBS and *α*B7-H3 monospecific CSAN controls against each individual data series by two-tailed unpaired t-test). **E**, CD4^+^ (left) and CD8^+^ (right) T cell populations were characterized at 24 and 72 hours after beginning co-culture by measuring CD45RO and CCR7 expression using flow cytometry. The phenotypes have been previously described as: naïve (CD45RO^−^/CCR7^+^), central memory (CD45RO^+^/CCR7^+^), effector memory (CD45RO^+^/CCR7^−^), effector (CD45RO^−^/CCR7^−^). (*P<0.05, ***P<0.001 with respect to *α*B7-H3-*α*CD3 bispecific CSAN treatment after 72 hours against each individual cell phenotype by two-tailed unpaired t-test). For every flow cytometry experiment, effector and target cells were determined by forward/side scatter and dead cells were removed using a viability dye (n = 9 using 3 PBMC donors total ± SEM). ELISA data was obtained using one PBMC donor but is representative of three different donors (**Supplementary Fig. S3 and S4**) (n = 3 wells per time point ± SD).

Memory T cell formation has been recognized as a vital component of a successful immune response to cancer for both CAR-T cells and targeted biologics (40). To investigate changes in T cell populations directed with controls or *α*B7-H3-*α*CD3 CSANs following 24 and 72 hour co-culture with a monolayer of ONS 2303 target cells, CD4^+^ and CD8^+^ T cell expression of memory cell surface markers, CD45RO and CCR7, were analyzed. Treatment with *α*B7-H3-*α*CD3 CSANs resulted in a significant increase in the observed percentage of central memory (CD45RO^+^/CCR7^+^) CD4^+^ T cells and a modest reduction in the amount of naïve (CD45RO^−^/CCR7^+^) CD4^+^ T cells after 72 hours of co-culture, relative to the PBS and monospecific CSAN controls as well as the bispecific CSAN treated 24 hour co-culture. (**Fig. 5E**). Similarly, at 72 hours, a significant increase in the amount of central memory CD8^+^ T cells was observed and, relative to the PBS and *α*B7-H3 monospecific controls, a significantly higher percentage of effector memory (CD45RO^+^/CCR7^−^) CD8^+^ T cells was also detected. A discernibly larger amount of effector (CD45RO^−^/CCR7^−^) CD8^+^ T cells directed with bispecific CSANs was observed after 24 hours of co-culture with ONS 2303 target cells and decreased over the next 48 hours. In contrast to the CD4^+^ T cells, a smaller percentage of naïve CD8^+^ T cells was observed as early as 24 hours and did not change over the course of the remaining 48 hours. These observed phenotypic changes did not significantly impact the relative total percentages of CD4^+^ and CD8^+^ T cells incubated with *α*B7-H3-*α*CD3 CSANs compared to those incubated with PBS and monospecific control CSANs (**Supplementary Fig. S5**).

### Bispecific CSAN activity is not dependent on MHC class I expression nor αB7-H3 avidity in vitro

To address the cytotoxic activity of the control *α*CD3 monospecific CSAN modified PBMCs observed during each live-cell imaging assay, we hypothesized that the MHC class I expressed on the target ONS 2303 cells is interacting with the unactivated PBMC T cell receptor (TCR) and, upon engagement with the *α*CD3 DHFR^2^ from the targeted or non-targeted CSANs, results in T cell activation and cytotoxicity. To test this hypothesis, the expression of MHC class I on the surface of ONS 2303 cells was knocked down by CRISPR/Cas9 targeting the β_2_-microglobulin gene, a critical component of MHC class I molecules. ONS 2303 cells that were double negative for MHC class I and β_2_-microglobulin proteins were collected by FACS and successful knock down (KD) was maintained for at least three culture passages after sorting, with only 1.5% of the cells shown to express MHC class I (**Supplementary Fig. S6A**). Cytotoxicity studies with wild type (WT) and KD ONS 2303 cells revealed that the loss of MHC class I expression minimally affected T cell mediated cytotoxicity directed by either *α*CD3 monospecific or *α*B7-H3-*α*CD3 bispecific CSANs (**Supplementary Fig. S6B-C**). Thus, both CSANs are postulated to direct T cell activity in a manner independent of MHC class I interactions on the target cells.

A second possibility for the non-targeted T cell cytotoxicity directed by the *α*CD3 monospecific CSANs is that the multivalency of the *α*CD3 scFv in the monospecific and bispecific CSANs has the potential to bind more than one T cell. Binding multiple T cells together could produce extracellular mechanical forces as well as CD3 antigen clustering and downstream signaling, all of which serve as a threshold for T cell activation (41). Additionally, UCHT1—the monoclonal antibody from which the *α*CD3 scFv is derived—is known to have mitogenic effects on T cells (42,43). This small amount of T cell activation/proliferation can induce cytokine release, particularly IFN*γ* secretion, which has been previously shown to increase MHC class II expression in the ONS76 target cells (44). Indeed, we have shown here that upon incubation with increasing IFN*γ* concentrations, the number of MHC class II antigens expressed on the surface of ONS 2303 cells increases accordingly (**Supplementary Fig. S7A-B**). Thus, we postulate that the non-targeted, multivalent *α*CD3 control CSANs elicit T cell cytotoxicity *in vitro* against ONS 2303 cells due to an increase in TCR:MHC class II interactions.

In previous publications, we have shown that the valency of each targeted (or non-targeted) monomer could be tuned by mixing different molar ratios of each monomer prior to addition of the bis-MTX dimerizer to form valency-modulated CSANs (27,30). Accordingly, we tested a few combinations of *α*CD3-valency-reduced targeted bispecific CSANs and the corresponding non-targeted monospecific CSANs (**Fig. 6A**). As previously observed, the equal-molar *α*CD3 monospecific CSANs, composed of an average ratio of 4 *α*CD3 DHFR^2^ and 4 non-targeted DHFR^2^ monomers, exhibited similar cytotoxicity to the corresponding (4:4) bispecific CSANs (**Fig. 6B**). However, when the valency of the *α*CD3 scFv is reduced, the non-targeted cytotoxic activity mediated by the control *α*CD3 monospecific CSANs is significantly diminished compared to the corresponding valency-tuned bispecific CSANs. Thus, tuning the valency of the *α*B7-H3-*α*CD3 CSANs to contain an average of only a single *α*CD3 moiety maintained a similar level of cytotoxic activity to that of the equal-molar (4:4) bispecific CSANs. Furthermore, the equal-molar (4:4) bispecific and monospecific CSANs equally elicited PBMC secretion of IFN*γ* and IL-2 after the 72 hour co-culture (**Fig. 6C**). Consistent with the observed reduction in cytotoxicity, the cytokine production from PBMCs modified with the *α*CD3 monospecific CSANs decreases incrementally in accordance with *α*CD3 scFv valency. Taken together, this data indicates that ONS 2303 medulloblastoma cells are particularly sensitive to non-targeted T cell activation elicited by multivalent *α*CD3 constructs. By taking advantage of the modular assembly of the CSANs, the potential for off-target T cell activation and cytotoxicity against ONS 2303 cells could be eliminated.

**Figure 6.**
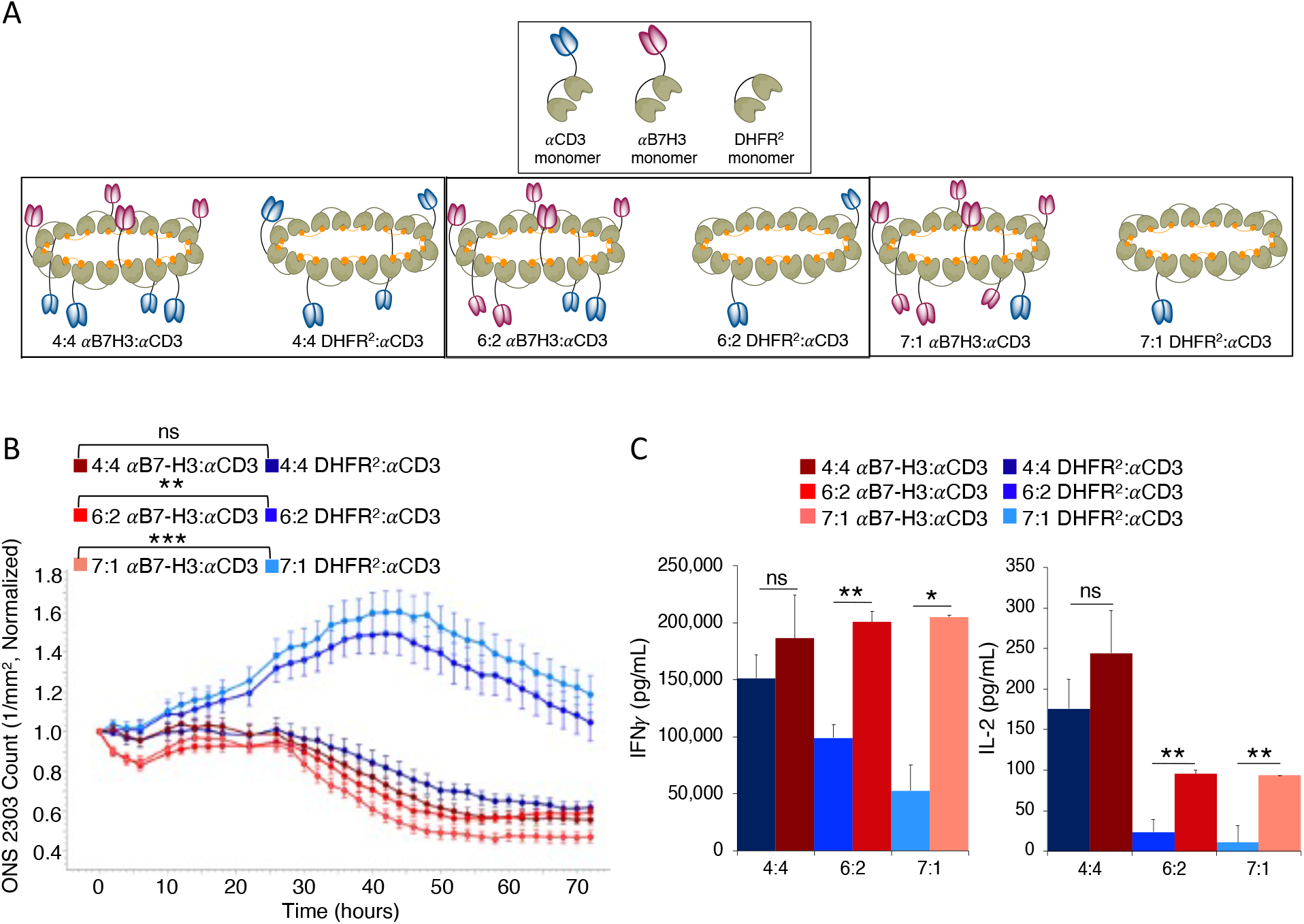
Tuning the valency of monospecific and bispecific CSANs controls non-targeted T cell activity. **A**, Reducing the valency of *α*CD3 monomers in the dimerized CSAN is done by increasing the molar ratios of either the *α*B7-H3 monomers or non-targeted DHFR^2^ monomers and decreasing the *α*CD3 monomers in the oligomerized CSAN. Shown is the average valency of the reduced *α*CD3 monomer CSANs. **B**, Target cells, unactivated PBMCs, and “valency-tuned” CSANs were co-cultured as previously described in a monolayer. ONS 2303 cell viability was monitored over a 72 hour period using an Incucyte SX5 at a fixed CSAN concentration (400nM) and PBMCs added at a 20:1 E:T ratio (Significance is indicated as **P<0.01, ***P<0.001 by two-tailed unpaired t-test between hours 14-72, n = 3 wells per time point ± SEM). **C**, Following the 72 hour viability study, the media from the co-culture was analyzed for IFN*γ* and IL-2 using a sandwich ELISA. (Significance is indicated as *P<0.05, **P<0.01 by two-tailed unpaired t-test, n = 3 wells per time point ± SD). Incucyte and ELISA data was obtained using one PBMC donor.

### Bispecific CSANs reduce tumor burden in orthotopic medulloblastoma models

To evaluate the *in vivo* anti-tumor activity of the *α*B7-H3-*α*CD3 CSAN directed T cells, we orthotopically injected 1×10^5^ ONS 2303 cells in the cerebellum of adult NRG mice. 11 days after tumor inoculation, the mice were separated into four groups and intravenously (IV) received 20×10^6^ unactivated PBMCs. Four days later, mice were intraperitoneally (IP) injected with the first dose of bispecific CSANs, monospecific CSANs, or PBS and received repeated booster injections every Monday, Wednesday, and Friday for 6 total treatments. 24 hours after the final treatment, mice were euthanized and their tissues collected for analysis (**Fig. 7A**). A significant reduction in the medulloblastoma tumor burden was observed in mice that received the bispecific CSANs compared to the PBS control 24 hours after the first treatment (**Fig. 7B-C**) while no significant weight loss was observed (**Supplementary Fig. S8**).

**Figure 7.**
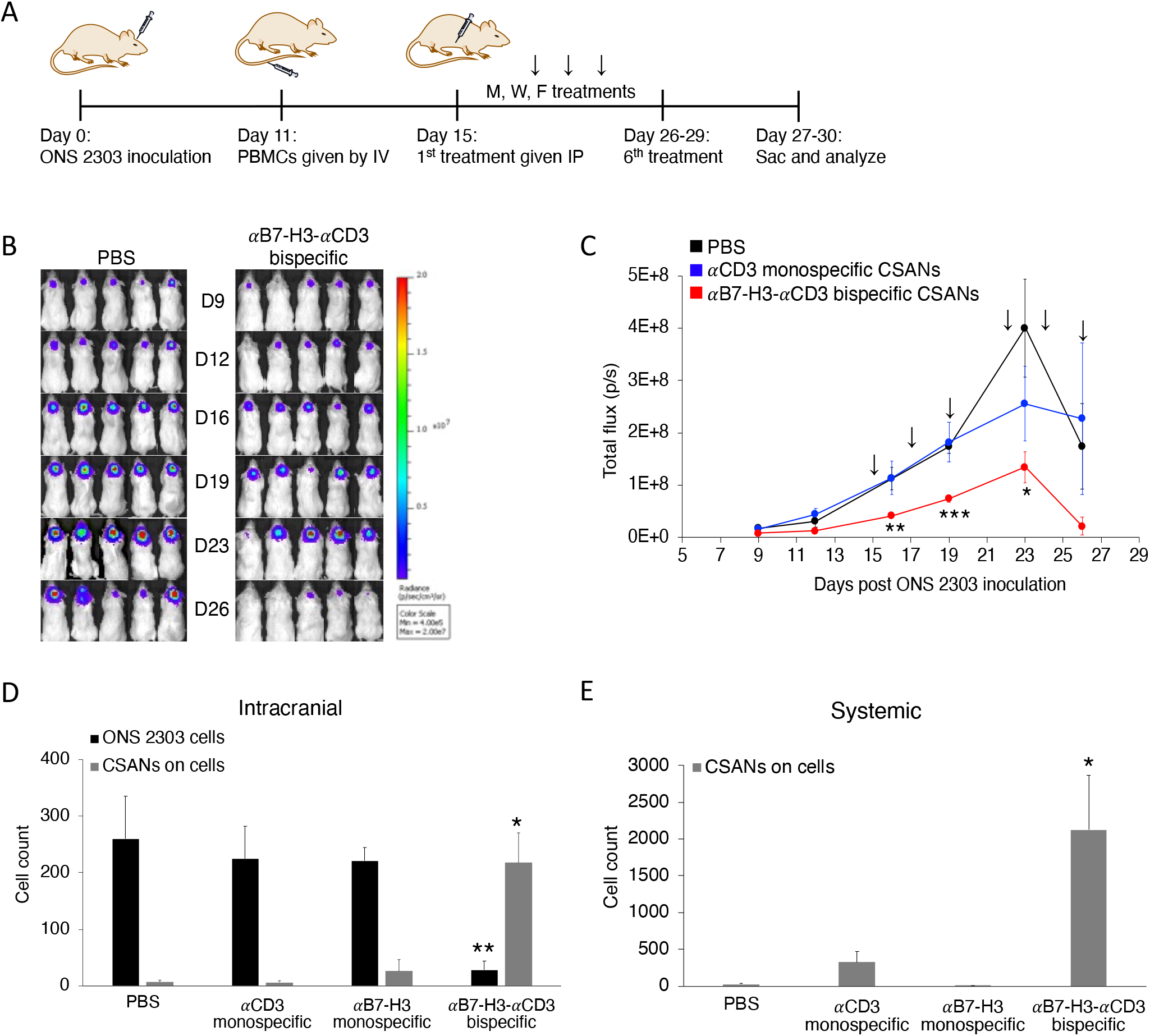
Systemically injected *α*B7-H3-*α*CD3 bispecific CSANs boost T cell protection for NRG mice against developing large orthotopic medulloblastoma tumors. **A**, Mouse model of orthotopic medulloblastoma. Intracranial injections were performed as previously described (34). 1×10^5^ ONS 2303 cells were intracranially injected into the cerebellum of adult NRG mice. Tumor engraftment was monitored by bioluminescent imaging and, 11 days after tumor inoculation, 20×10^6^ unactivated human PBMCs were IV injected. 4 days after PBMCs were given, mice received 1mg/kg CSAN treatments or PBS control IP injections. Booster injections were given every Monday, Wednesday, and Friday for 6 total treatments; the 6^th^ treatment was staggered for some mice so that the flow cytometry analysis on live cells would happen about 24 hours after the final injection. **B**, Bioluminescent IVIS images of ONS 2303 tumors. **C**, Tumor progression was monitored by bioluminescent photometry where the average flux values (p/s, calculated by Living Image software) are shown for treatment and control groups (*P<0.05, **P<0.01, ***P<0.001 with respect to the PBS control by two-tailed unpaired t-test). **D**, A piece of the cerebellum that had been inoculated was removed and dissociated into a single cell suspension to enable measurements of viable ONS 2303 (GFP^+^ expression) and CSAN (Flag tag) presence on target brain and effector T cells by flow cytometry (**P<0.01, ***P<0.001 with respect to PBS and both monospecific control CSANs against each individual data series by two-tailed unpaired t-test). **E**, Blood was collected at the time of sacrifice and red blood cells were removed using ACK buffer. The remaining cell populations were analyzed for CSAN (Flag tag) presence by flow cytometry (*P<0.05 with respect to PBS and both monospecific control CSANs by two-tailed unpaired t-test). For every flow cytometry experiment, effector and target cells were determined by forward/side scatter and dead cells were removed using a viability dye (n = 5-7 mice per treatment group, data is displayed as mean ± SEM).

To demonstrate that the bispecific CSANs were able to effectively cross the BTB to direct the PBMC anti-tumor response, we quantified the nanorings bound to remaining intracranial effector/target cells and compared that to systemically circulating CSANs. Mice treated with the bispecific CSANs had significantly fewer (8-fold less) ONS 2303 target cells remaining within the intracranial samples correlating well with the significantly higher amount of bispecific nanorings (8.2-fold more) detected on the target and effector cells compared to those treated with the monospecific CSANs and PBS (**Fig. 7D**). Additionally, significant quantities of bispecific CSANs were detected bound to systemic lymphocytes compared to the non-targeted controls (**Fig. 7E**). The small amount of *α*CD3 monospecific CSANs circulating systemically has been observed previously where a significant proportion of these non-targeted nanorings were accumulated within the major clearance organs 24 hours post-injection (26).

### Bispecific CSANs selectively activate and encourage T cell expansion and memory formation in vivo

Successful T cell engraftment and expansion is a hallmark of an efficacious response to targeted cancer therapies (45). Here, we evaluated this response in the orthotopic medulloblastoma model using *α*B7-H3-*α*CD3 CSANs as previously discussed. Unlike the *in vitro* results (**Supplementary Fig. S5**), repeated treatments of bispecific CSANs significantly impacted CD4^+^ and CD8^+^ T cell expansion both intracranially and systemically (**Fig. 8A-B**). Interestingly, the actual values between the CD4^+^ and CD8^+^ T cells were similar for each sample type, which likely affected the anti-tumor response observed above as it has been previously demonstrated that expansion of both CD4^+^ and CD8^+^ T cells is correlated with prolonged survival in cancer patients (46,47).

**Figure 8.**
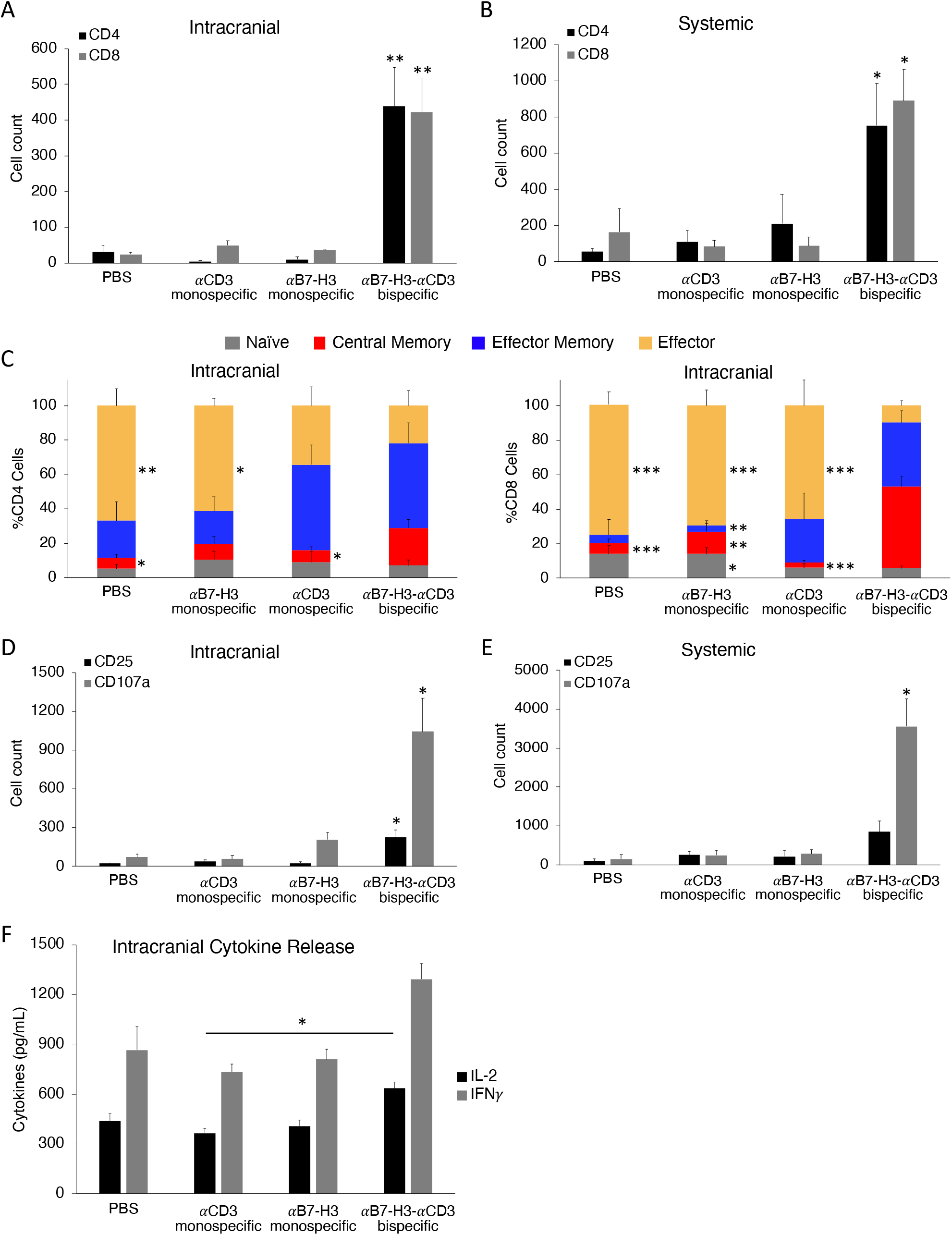
*α*B7-H3-*α*CD3 bispecific CSANs selectively activate and encourage CD4^+^ and CD8^+^ T cell expansion and central memory T cell formation *in vivo*. The expansion of CD4^+^ and CD8^+^ T cells was measured both **A**, intracranially using the cerebellum single cell suspension; and **B**, systemically using the blood collected from each mouse (*P<0.05, **P<0.01 with respect to PBS and *α*CD3 monospecific CSAN control against each individual data series by two-tailed unpaired t-test). **C**, Intracranial CD4^+^ (left) and CD8^+^ (right) T cell populations were characterized using the cerebellum single cell suspension and measuring CD45RO and CCR7 expression by flow cytometry. The phenotypes have been previously described as: naïve (CD45RO^−^/CCR7^+^), central memory (CD45RO^+^/CCR7^+^), effector memory (CD45RO^+^/CCR7^−^), and effector (CD45RO^−^/CCR7^−^). (*P<0.05, **P<0.01, ***P<0.001 with respect to *α*B7-H3-*α*CD3 bispecific CSAN treatment against each individual cell phenotype by two-tailed unpaired t-test). T cell expression of activation markers, CD25 and CD107a, were measured by flow cytometry using **D**, the cerebellum single cell suspension and **E**, the lymphocytes collected from the peripheral blood of each mouse (*P<0.05 with respect to PBS and both monospecific CSAN controls against each individual data series by two-tailed unpaired t-test). **F**, A separate piece of the cerebellum that had been inoculated with ONS 2303 cells was collected and flash frozen before lysis. The cerebellar lysate was used to measure the intracranial T cell secretion of IFN*γ* and IL-2 by sandwich ELISA (*P<0.05 by two-tailed unpaired t-test). For every flow cytometry experiment, effector and target cells were determined by forward/side scatter and dead cells were removed using a viability dye. (n = 5-7 mice per treatment group, data is displayed as mean ± SEM).

Multiple groups have suggested that a persistent memory T cell population is necessary for a complete response to adoptive immunotherapy and a few have narrowed this even further proposing that the central memory phenotype is optimal for a complete and sustained response (48,49). We demonstrated that mice treated with the bispecific CSANs showed significant expansion of both CD4^+^ and CD8^+^ central memory T cells intracranially (3.5-fold and 7.8-fold, respectively compared to the PBS control) and significantly diminished percentages of effector T cells (**Fig. 8C**), which is in accordance with the well documented life cycle and differentiation of antigen-specific T cells (40). Effector and central memory T cell phenotypes were also present systemically, but minimal differences were observed between the treated and control groups (**Supplementary Fig. S9**).

Due to the successful engraftment and expansion of PBMCs within our medulloblastoma mouse model, the resultant degree of *in vivo* T cell activation was assessed by quantifying CD25 and CD107a expression. Both activation markers were elevated intracranially for mice treated with the *α*B7-H3-*α*CD3 CSANs indicating that booster injections can sustain T cell activation across the BTB, intratumorally (**Fig. 8D**). Systemic lymphocytes from the same group of mice, however, did not express a significant amount of late activation marker CD25, but did exhibit increased quantities of the degranulation marker, CD107a (**Fig. 8E**). When co-cultured with medulloblastoma cell line, ONS 2303, *α*B7-H3-*α*CD3 CSAN modified PBMCs increased secretion of IFN*γ* and IL-2 (**Fig. 5A-B**). To test whether bispecific CSANs similarly affect T cell cytokine secretion *in vivo*, we analyzed cytokine concentrations within frozen sections of intracranial medulloblastoma tumors from the orthotopic NRG mouse model. Cytokine analysis revealed that, while the bispecific CSAN treated mice did exhibit increased concentrations of IFN*γ* and IL-2 intracranially, only IL-2 production was significantly higher compared to the *α*CD3 monospecific control (**Fig. 8F**). Low levels of cytokine production within medulloblastoma tumors has been well documented (50) and could actually prove to be protective as cytokine storm occurring within a pediatric brain tumor could be life-threatening (51).

Many different adoptive immunotherapy approaches have had to overcome suppressive tumor microenvironments that can lead to immune escape, particularly by managing regulatory T cell (Treg) activity and blocking immunosuppressive checkpoint molecule interactions (52,53). To determine whether the bispecific CSAN treatments affected the formation and infiltration of Treg cell populations, lymphocytes remaining within tumor tissues from the orthotopic study expressing CD4^+^/CD25^+^/FOXP3^+^ were quantified. In general, very few Treg cells were present intratumorally with significantly less discovered in the brains of mice treated with bispecific CSANs (**Supplementary Fig. S10A**). Additionally, we sought to understand whether *α*B7-H3-*α*CD3 CSAN therapy played a role in cancer cell and immune cell upregulation of checkpoint molecule expression intratumorally. CTLA-4 is an inhibitory receptor expressed by activated T cells and blockade of its inhibitory signaling has been shown to restore anti-tumor immunity and promote tumor regression (53,54). We found that the number of cells expressing CTLA-4 was about 8-fold higher intracranially compared to the indicated controls but was not significant systemically (**Supplementary Fig. S10B**). It is known that CTLA-4 expression is constitutive on Treg cells but is also upregulated in normal T cells upon activation (55). Due to the sparse population of Treg cells found intracranially, these results indicate that CTLA-4 may be overexpressed by the activated T cells directed with bispecific CSANs and represents the potential for complementary checkpoint therapy. Consistent with previous reports of limited PD-L1 expression on medulloblastoma cells (56), we were unable to draw conclusions from this study on the effect of bispecific CSAN therapy on PD-1 and PD-L1 expression (data not included).

## Conclusions

Here we have investigated the feasibility of targeting B7-H3, a cancer antigen expressed on a wide variety of pediatric solid tumors, using a non-genetic approach to direct cell-to-cell interactions with CSANs. We demonstrated that the *α*B7-H3-*α*CD3 CSANs directed potent and specific T cell binding and activity *in vitro* against a cultured monolayer of medulloblastoma cells and increased the T cell infiltration and cytotoxicity of medulloblastoma spheroids. It was also confirmed that CSAN directed T cells are activated without TCR:MHC class I interactions, thus loss of MHC class I expression as an immune evasion mechanism will not affect bispecific CSAN efficacy. Importantly, the non-targeted cytotoxicity caused by T cells directed with the multivalent *α*CD3 monospecific control CSANs was reduced by tuning the valency of the *α*CD3 scFv in the nanoring scaffold; but this reduction in *α*CD3 valency did not affect the activity of the targeted *α*B7-H3-*α*CD3 CSANs. We hypothesize that the multivalency and mitogenicity of the *α*CD3 scFv could trigger T cell activation and secretion of IFN*γ*, causing an upregulation of MHC class II expression on the ONS 2303 cells, resulting in non-targeted T cell cytotoxicity due to an increase in TCR:MHC class II interactions. Interestingly, this non-targeted response directed by the *α*CD3 monospecific CSANs is not observed *in vivo*.

Understanding the magnitude of T cell activation generated by bispecific CSAN targeting and characterizing the resultant immunosuppressive tumor microenvironment are all critical aspects to consider when designing a brain tissue targeting immunotherapeutic. Therefore, we reveal that targeted bispecific CSANs significantly increase T cell secretion of stimulating cytokines and expression of hallmark T cell activation receptors both *in vitro* and *in vivo*. Additionally, we demonstrated that *in vivo* IP injections of *α*B7-H3-*α*CD3 CSANs were able to traverse the peritoneum and cross the BTB to direct selective T cell responses within intracranial brain tumors. However, because the goal of this study was to investigate the intratumoral CSAN directed T cell response, we were unable to determine if complete eradication of the medulloblastoma tumors in NRG mice was achieved. On going studies suggest that tumor eradiation could be achieved with a longer treatment regimen and will be reported in due course. Additionally, mice treated with the bispecific CSANs elicited robust CD8^+^ central memory T cell formation within the tumor itself, which has been associated with better prognosis and more potent anti-tumor responses in cancer patients (45) but has yet to be elucidated for B7-H3 CAR-Ts (13). Using this model, we were also able to determine the degree of Treg response as well as characterize the immune checkpoint molecule expression intracranially which could be further explored in combination immunotherapies.

Although many preclinical studies have identified B7-H3 overexpression in solid tumors, it is maintained that expression can be detected in normal tissues, albeit in restricted quantities (13–15). To address off-targeting concerns, we derived our bispecific nanorings from the 8H9 antibody which has been safely used as a radioimmunotherapy in pediatric patients and has been incorporated as a chimeric antigen receptor into natural killer cells with positive outcomes (19,22,57). Throughout this study, typical on-target off-tumor toxicities were not observed, such as loss of body weight or neurotoxicity associated with targeting B7-H3. However, the safety of the *α*B7-H3-*α*CD3 CSANs can only be accurately investigated in a well-designed immunocompetent model as the humanized 8H9 scFv does not cross react with murine B7-H3 antigen (31). Despite this, if toxicities should arise, the CSANs can be quickly disassembled upon injection of FDA-approved drug trimethoprim, preventing T cell overstimulation (26). Additionally, we found here and in previous reports that tuning the valency of the monomers affects the overall CSAN affinity and off-target T cell activity (27,30).

In summary, we engineered, characterized, and tested an *α*B7-H3-*α*CD3 bispecific CSAN that effectively crossed the BTB and directed a potent anti-tumor T cell response against established medulloblastoma tumors. *α*B7-H3-*α*CD3 CSANs did not cause evident toxicity, but rather encouraged T cell expansion and memory cell formation providing evidence of its potential as a new and complementary therapeutic option for pediatric solid tumors.

## Acknowledgements

The authors would like to thank Dr. Benjamin Hackel for use of the MS1_hB7-H3_ cell line, Wendy Hudson for breeding and supplying NRG animals, and the University Imaging Centers and University Flow Cytometry Resource at the University of Minnesota for experimental and technical support. EA Mews is supported by the National Science Foundation GRFP Fellowship. Support of this study by NCI R01CA247681(CRW) is gratefully acknowledged.

