## Supplemental Info for "Multivalent, Bispecific *α*B7-H3-*α*CD3 Chemically Self-Assembled Nanorings Direct Potent T-cell Responses Against Medulloblastoma"

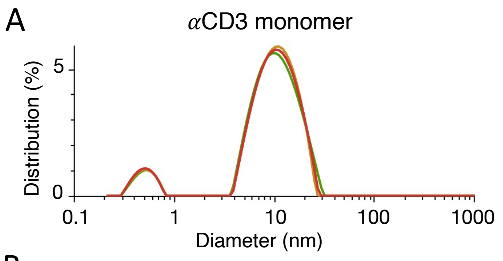


B

| | Mean Diameter (nm) | Polydispersity (%) |
|---|---|---|
| αCD3 monomer | 9.2 ± 0.3 | 24 ± 1 |
| αB7-H3 monomer | 9.4 ± 0.2 | 16 ± 2 |
| αB7-H3 monospecific CSAN | 15.8 ± 0.7 | 22 ± 3 |

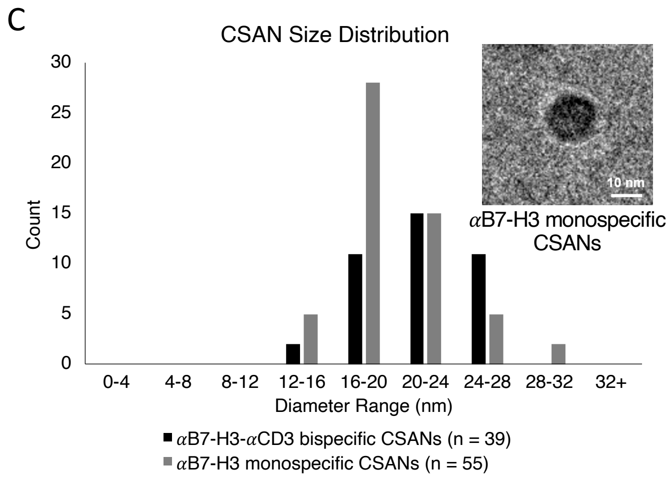


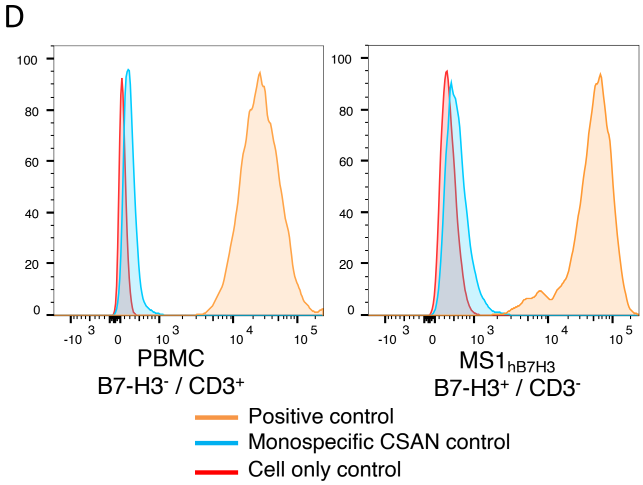


### Supplementary Figure S1.

αB7-H3 CSAN characterization. **A,** The hydrodynamic diameter and polydispersity index (PDI) of the αCD3 monomer was determined using dynamic light scattering (DLS); representative image of 2 experiments shown. **B,** Table summarizing the hydrodynamic diameters and polydispersity indexes of the tested monomers and CSANs from A, and Fig. 2C. **C,** Cryo-transmission electron microscopy (cryo-TEM) size distribution analysis comparing bispecific (average = 22 ± 3 nm) and monospecific αB7-H3 CSANs (average = 20 ± 3 nm) and a representative image of αB7-H3 monospecific CSANs. Scale bar, 10nm. **D,** Monospecific and positive controls from the CSAN bispecificity binding study in Fig. 2E.

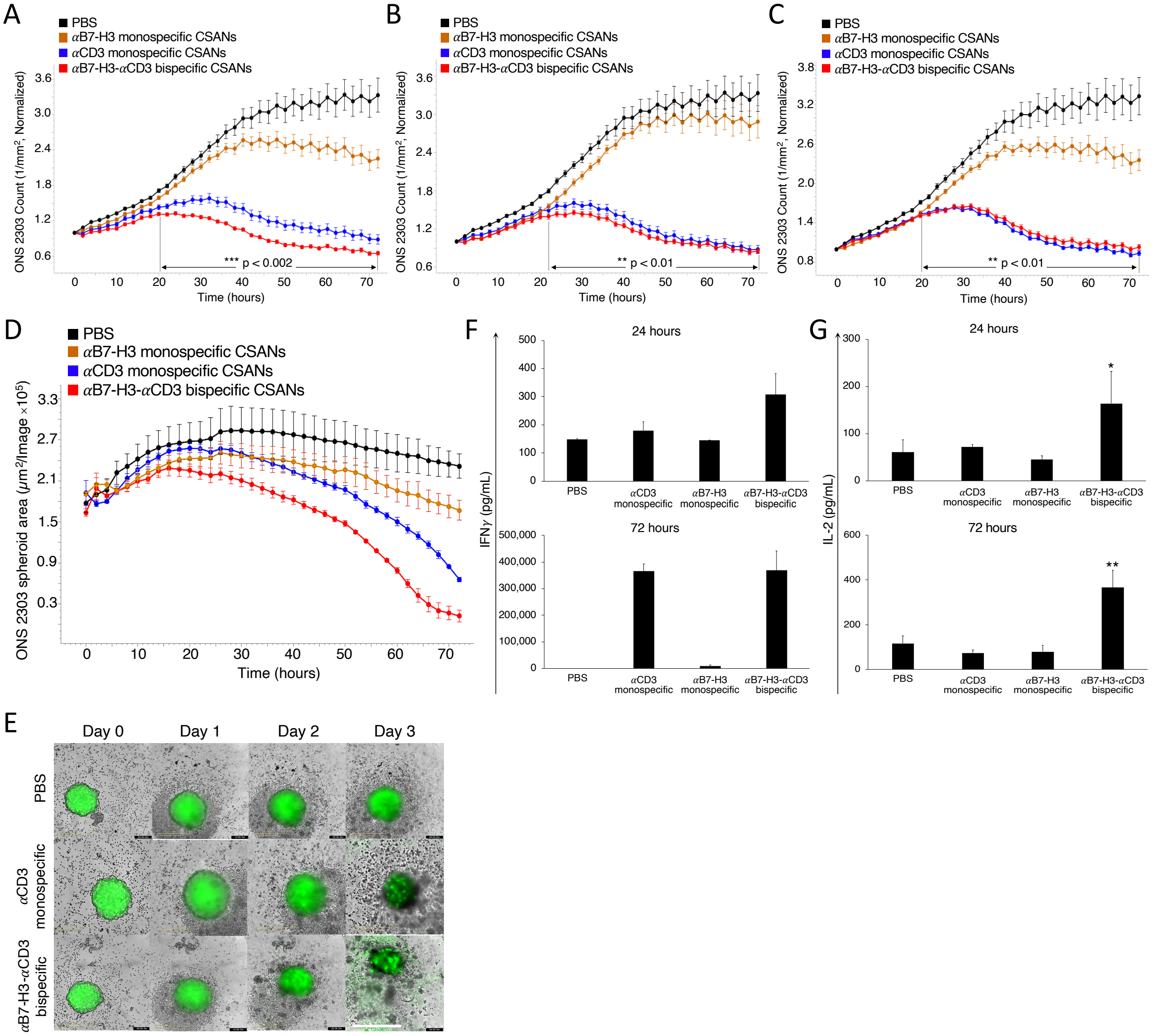


### Supplementary Figure S2.

Donor #1: αB7-H3-αCD3 CSANs at lower concentrations enable selective T cell activation and cytotoxicity against B7-H3$^+$ target cells. ONS 2303 was seeded into a 96 well plate and cultured as a monolayer. Unactivated PBMCs were then added to the target cells 16 hours later with either monospecific or bispecific CSANs all at once. ONS 2303 cell viability was monitored over a 72 hour period using an Incucyte SX5 at different CSAN concentrations: **A,** 200nM, **B,** 100nM, **C,** 50nM and fixed E:T ratio (20:1) (**P<0.01, ***P<0.002 with respect to PBS control by two-tailed unpaired t-test). **D,** ONS 2303 was seeded into a 96 well ULA plate and cultured as a spheroid for 2 days. Unactivated PBMCs were then added to the spheroids with either monospecific or bispecific CSANs all at once. The spheroid viability was monitored over a 72 hour period using an Incucyte SX5 at a fixed CSAN concentration (200nM) and fixed E:T ratio (5:1) (**P<0.01 with respect to PBS control by two-tailed unpaired t-test). **E,** Representative images of ONS 2303 spheroids from **D** treated with either PBS, αCD3 monospecific, or αB7-H3-αCD3 CSANs over time. Scale bar, 400µm. Following the viability study, at the specified time points, the media from the 200nM co-culture was analyzed for **F,** IFNγ and **G,** IL-2 using a sandwich ELISA (*P<0.05, **P<0.01 with respect to no treatment and monospecific control CSANs by two-tailed unpaired t-test). Incucyte data (n = 3 wells per time point ± SEM) and ELISA data (n = 3 wells per time point ± SD). All Incucyte images were taken with a 10x objective. Normalized data is shown as a ratio of the ONS 2303 viability compared to viability at time zero.

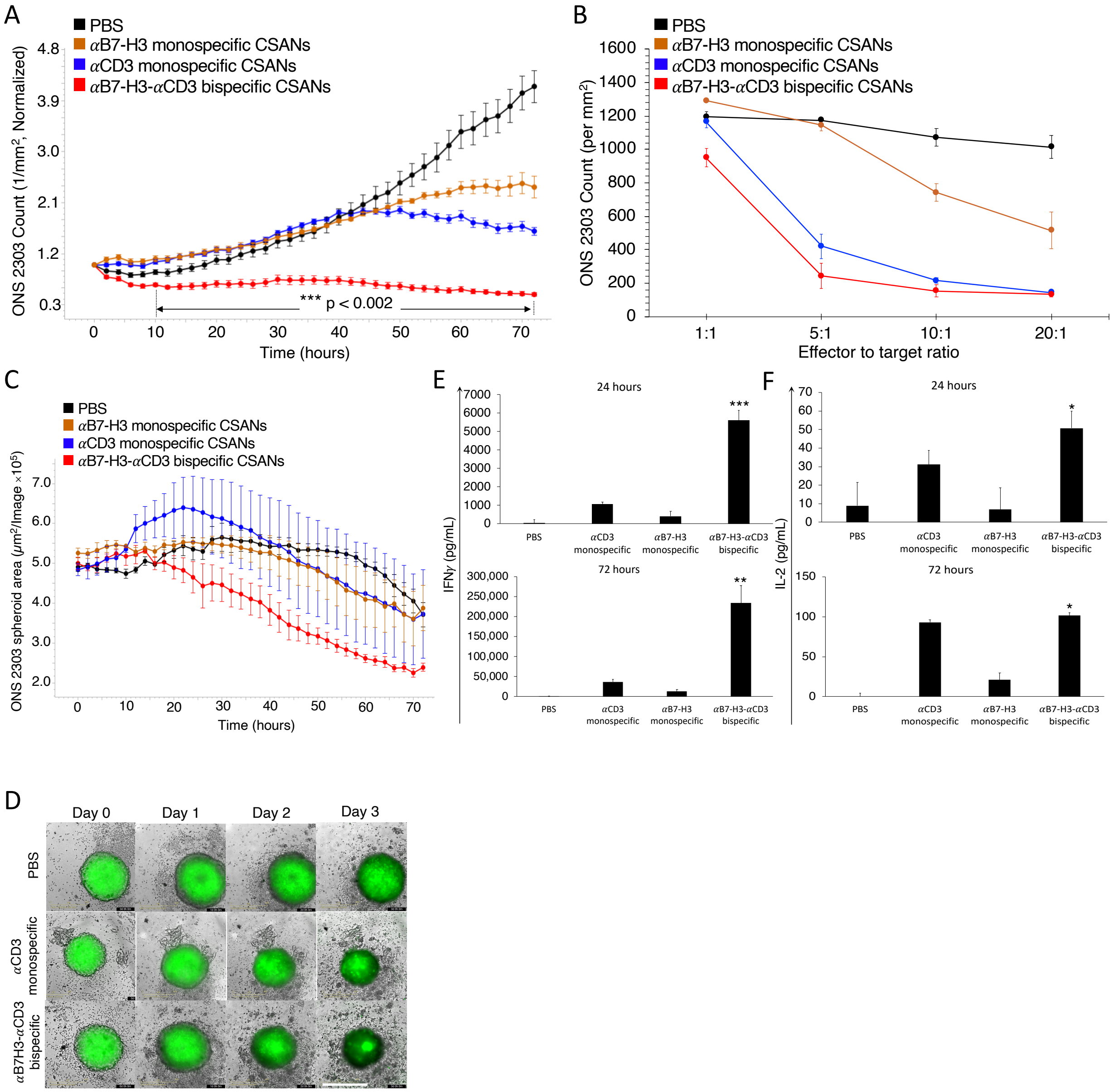


### Supplementary Figure S3.

Donor #2: αB7-H3-αCD3 CSANs enable selective T cell activation and cytotoxicity against B7-H3$^+$ target cells. ONS 2303 was seeded into a 96 well plate and cultured as a monolayer. Unactivated PBMCs were then added to the target cells 16 hours later with either monospecific or bispecific CSANs all at once. **A,** ONS 2303 cell viability was monitored over a 72 hour period using an Incucyte SX5 at a fixed CSAN concentration (400nM) and fixed E:T ratio (20:1) (***P<0.002 with respect to PBS control by two-tailed unpaired t-test). **B,** Different E:T ratios were similarly analyzed at a fixed CSAN concentration (400nM) where each bullet point represents the final ONS 2303 viable cell count at the end of the 72 hour cytotoxicity study (*P<0.05, ***P<0.001 with respect to PBS control by two-tailed unpaired t-test). **C,** ONS 2303 was seeded into a 96 well ULA plate and cultured as a spheroid for 2 days. Unactivated PBMCs were then added to the spheroids with either monospecific or bispecific CSANs all at once. The spheroid viability was monitored over a 72 hour period using an Incucyte SX5 at a fixed CSAN concentration (400nM) and fixed E:T ratio (5:1) (**P<0.01 with respect to PBS control by two-tailed unpaired t-test).

**D,** Representative images of ONS 2303 spheroids from **C** treated with either PBS, $\alpha$CD3 monospecific, or $\alpha$B7-H3-$\alpha$CD3 CSANs over time. Scale bar, 400µm. Following the viability study, at the specified time points, the media from the co-culture was analyzed for **E,** IFN$\gamma$ and **F,** IL-2 using a sandwich ELISA (*P<0.05, **P<0.01, ***P<0.001 with respect to no treatment and monospecific control CSANs by two-tailed unpaired t-test). Incucyte data (n = 3 wells per time point ± SEM) and ELISA data (n = 3 wells per time point ± SD) was obtained using one PBMC donor but is representative of three different donors (**Figures 4 and S4**). All Incucyte images were taken with a 10x objective. Normalized data is shown as a ratio of the ONS 2303 viability compared to viability at time zero.

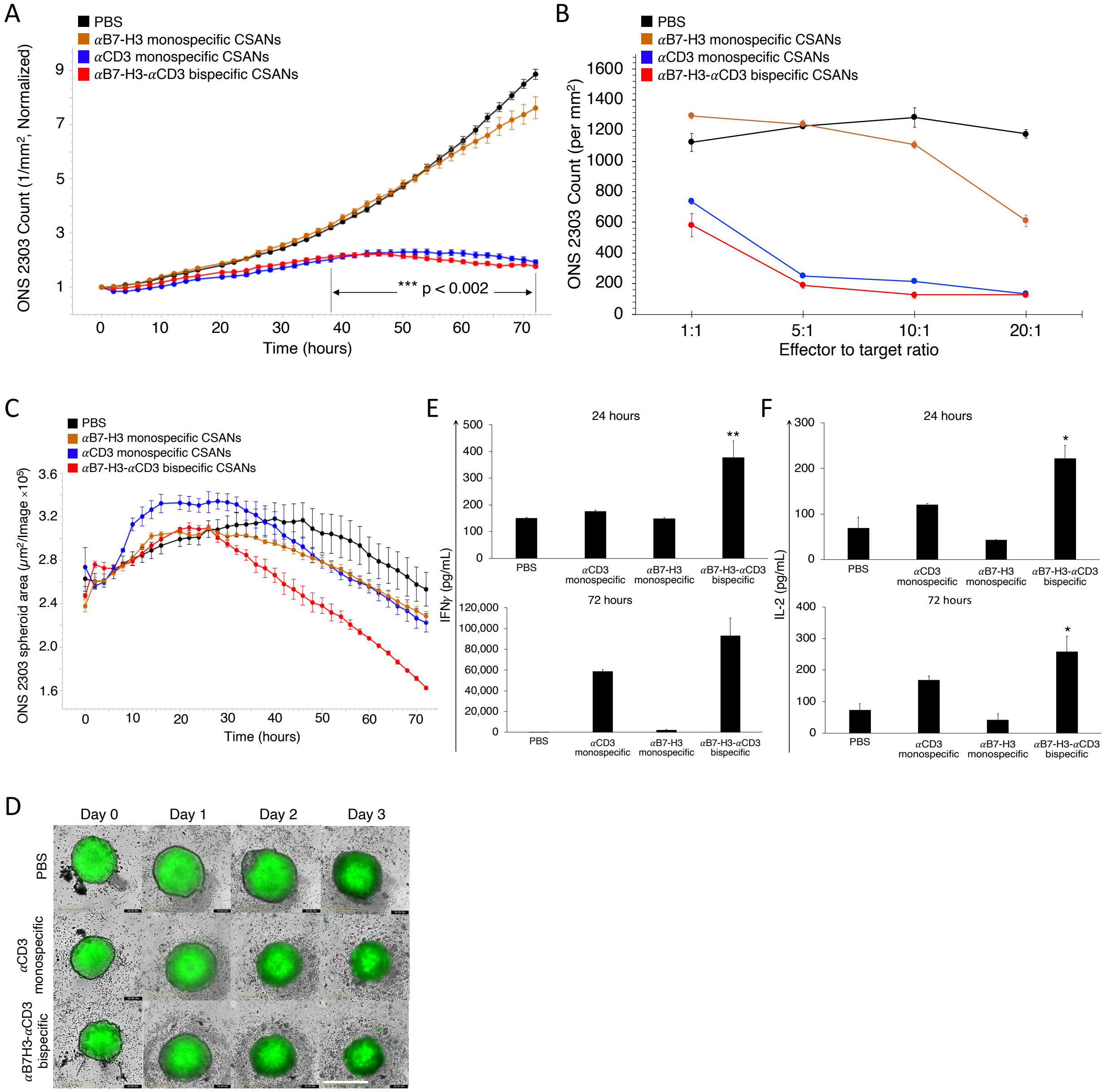


### Supplementary Figure S4.

Donor #3: αB7-H3-αCD3 CSANs enable selective T cell activation and cytotoxicity against B7-H3$^{+}$ target cells. ONS 2303 was seeded into a 96 well plate and cultured as a monolayer. Unactivated PBMCs were then added to the target cells 16 hours later with either monospecific or bispecific CSANs all at once. **A,** ONS 2303 cell viability was monitored over a 72 hour period using an Incucyte SX5 at a fixed CSAN concentration (400nM) and fixed E:T ratio (20:1) (***P<0.002 with respect to PBS control by two-tailed unpaired t-test). **B,** Different E:T ratios were similarly analyzed at a fixed CSAN concentration (400nM) where each bullet point represents the final ONS 2303 viable cell count at the end of the 72 hour cytotoxicity study (*P<0.05, ***P<0.001 with respect to PBS control by two-tailed unpaired t-test). **C,** ONS 2303 was seeded into a 96 well ULA plate and cultured as a spheroid for 2 days. Unactivated PBMCs were then added to the spheroids with either monospecific or bispecific CSANs all at once. The spheroid viability was monitored over a 72 hour period using an Incucyte SX5 at a fixed CSAN concentration (400nM) and fixed E:T ratio (5:1) (**P<0.01 with respect to PBS control by two-tailed unpaired t-test). **D,** Representative images of ONS 2303 spheroids from **C** treated with either PBS, αCD3 monospecific, or αB7-H3-

$\alpha$CD3 CSANs over time. Scale bar, 400µm. Following the viability study, at the specified time points, the media from the co-culture was analyzed for **E,** IFN$\gamma$ and **F,** IL-2 using a sandwich ELISA (*P<0.05, **P<0.01 with respect to no treatment and monospecific control CSANs by two-tailed unpaired t-test). Incucyte data (n = 3 wells per time point ± SEM) and ELISA data (n = 3 wells per time point ± SD) was obtained using one PBMC donor but is representative of three different donors (**Figures 4 and S3**). All Incucyte images were taken with a 10x objective. Normalized data is shown as a ratio of the ONS 2303 viability compared to viability at time zero.

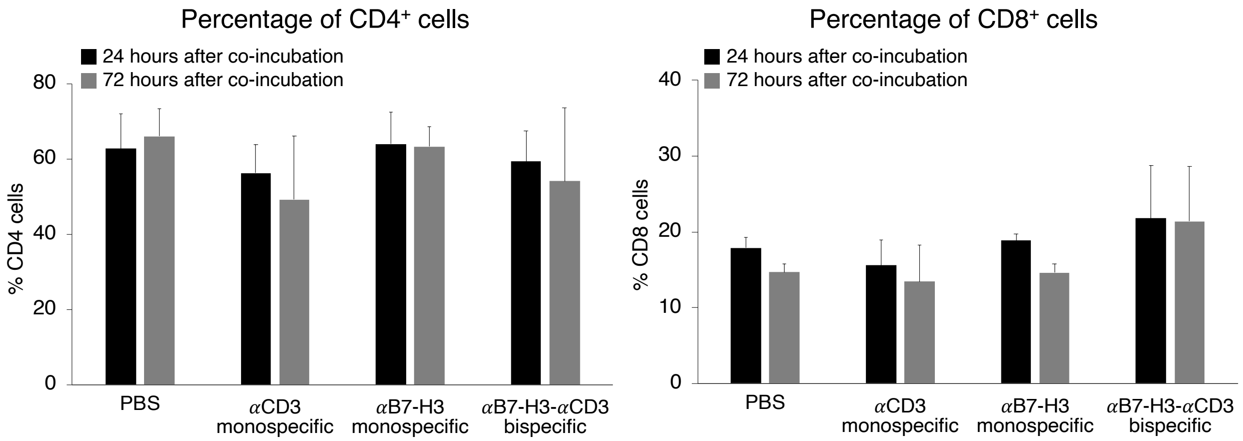


### Supplementary Figure S5.

Monitoring $CD4^+$ and $CD8^+$ T cell expansion in vitro by flow cytometry. ONS 2303 was seeded into a 96 well plate and cultured as a monolayer. Unactivated PBMCs were then added to the target cells 16 hours later with either monospecific or bispecific CSANs all at once. The expansion of $CD4^+$ and $CD8^+$ T cells was measured at 24 and 72 hours after beginning the co-culture. Effector cells were differentiated by forward/side scatter and dead cells were removed using a viability dye (n = 9 using 3 PBMC donors total, data is displayed as mean ± SEM).

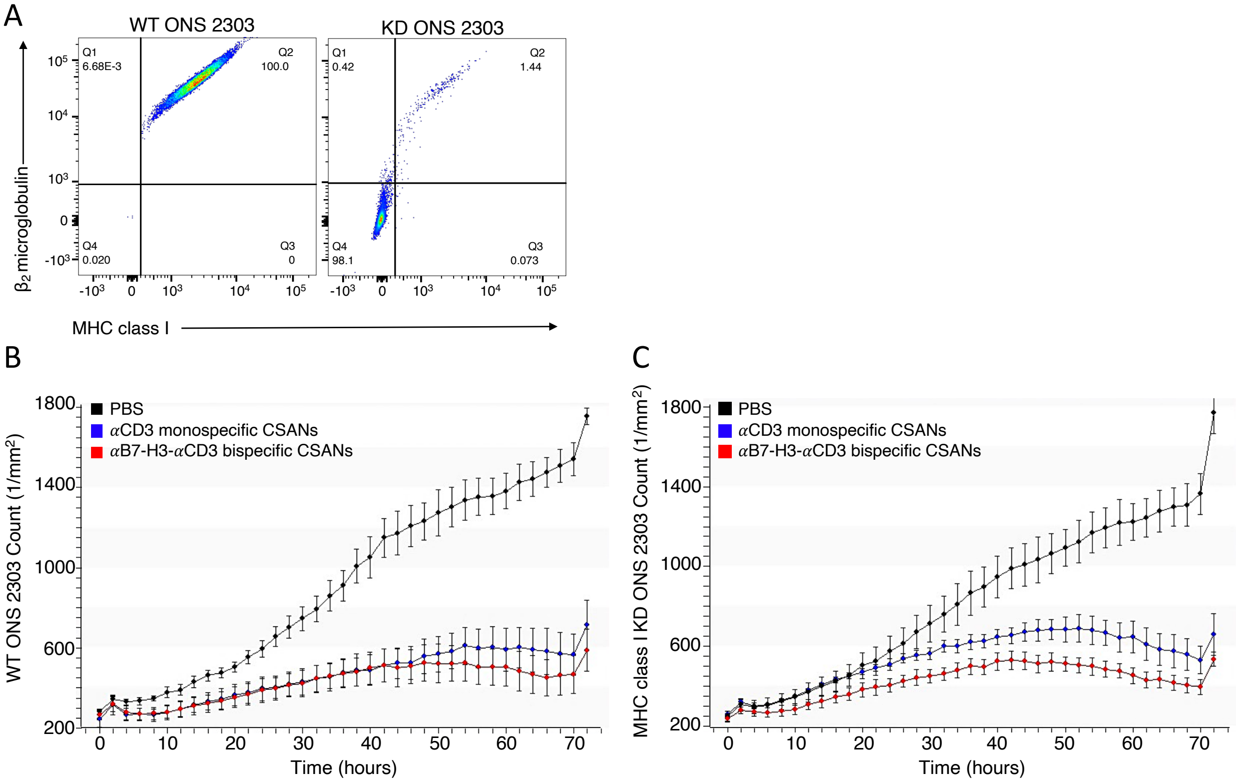


### Supplementary Figure S6.

Knocking down MHC class 1 expression on ONS 2303 cells does not inhibit $\alpha$B7-H3-$\alpha$CD3 CSAN mediated T cell activity nor does it reduce $\alpha$CD3 monospecific CSAN mediated T cell activity. $\beta_2$ microglobulin (B2M), a critical component of MHC class I molecules, was knocked down (KD) using CRISPR and the resultant MHC class I KD ONS 2303 cells were FACS sorted. Cells that were double negative for MHC class I and B2M were collected and expanded. **A,** Post FACS sort results of wild type (WT) and expanded MHC class I KD ONS 2303 cells where only ~1% of cells maintain MHC class I expression. Target cells, unactivated PBMCs, and CSANs were co-cultured as previously described. **B,** WT ONS 2303 and **C,** MHC class I KD ONS 2303 cell viability was compared over a 72 hour period using an Incucyte Zoom at a fixed CSAN concentration (400nM) and fixed E:T ratio (20:1). Incucyte images were taken with a 10x objective (n = 3 wells per time point ± SEM).

A

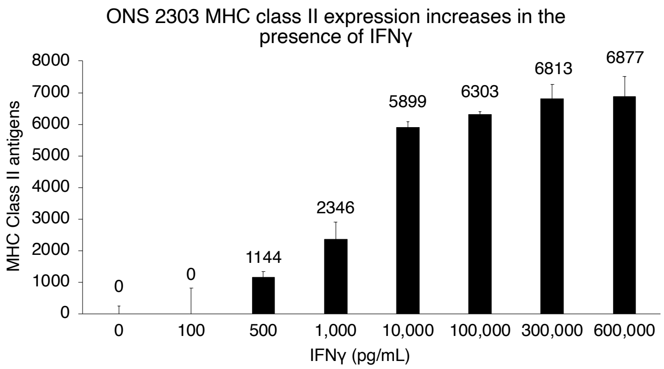


B

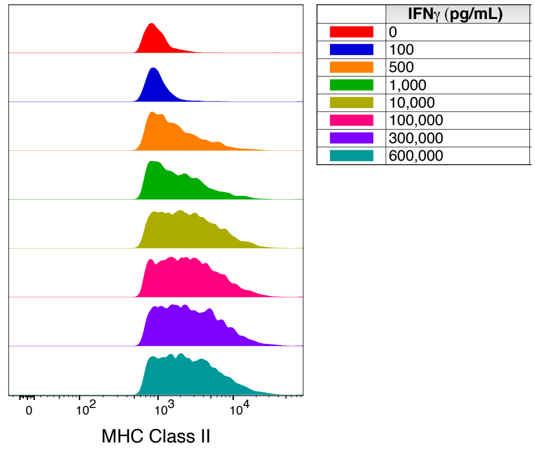


### Supplementary Figure S7

ONS 2303 expression of MHC class II antigens increases in the presence of IFN$\gamma$. **A)** Quantification of MHC class II antigen expression on the surface of ONS 2303 target cells cultured with increasing concentrations of IFN$\gamma$ for 20 hours. (Data is displayed as mean ± SD, n = 3 technical replicates). **B)** Representative flow cytometry curves used to quantify MHC class II expression in A.

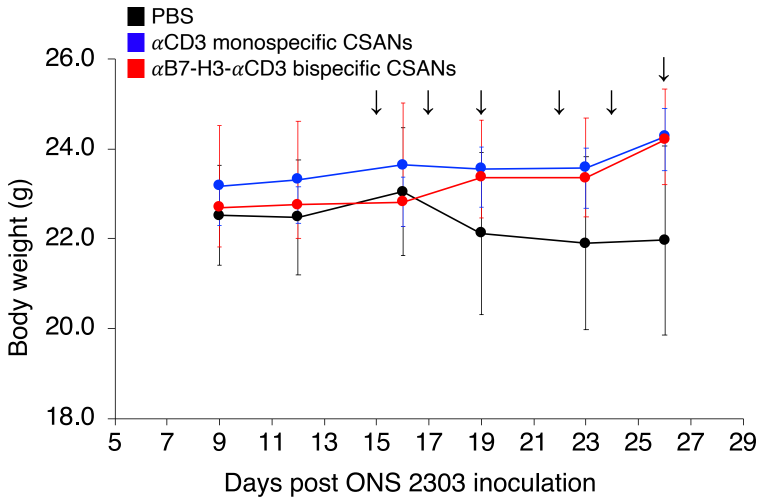


### Supplementary Figure S8.

Systemically injected αB7-H3-αCD3 CSANs boost T cell protection for NRG mice against developing large orthotopic medulloblastoma tumors. Intracranial injections were performed as previously described (34). Briefly, $1\times10^5$ ONS 2303 cells were intracranially injected into the cerebellum of adult NRG mice. Tumor engraftment was monitored by bioluminescent imaging and, 11 days after tumor inoculation, $20\times10^6$ unactivated human PBMCs were IV injected. 4 days after PBMCs were given, mice received 1mg/kg CSAN treatments or PBS control IP injections. Booster injections were given every Monday, Wednesday, and Friday for 6 total treatments; the 6$^{th}$ treatment was staggered for some mice so that the flow cytometry analysis on live cells would happen about 24 hours after the final injection. Average body weight was monitored and recorded for each treatment group. Arrows indicate treatment days (n = 5-7 mice per group ± SEM).

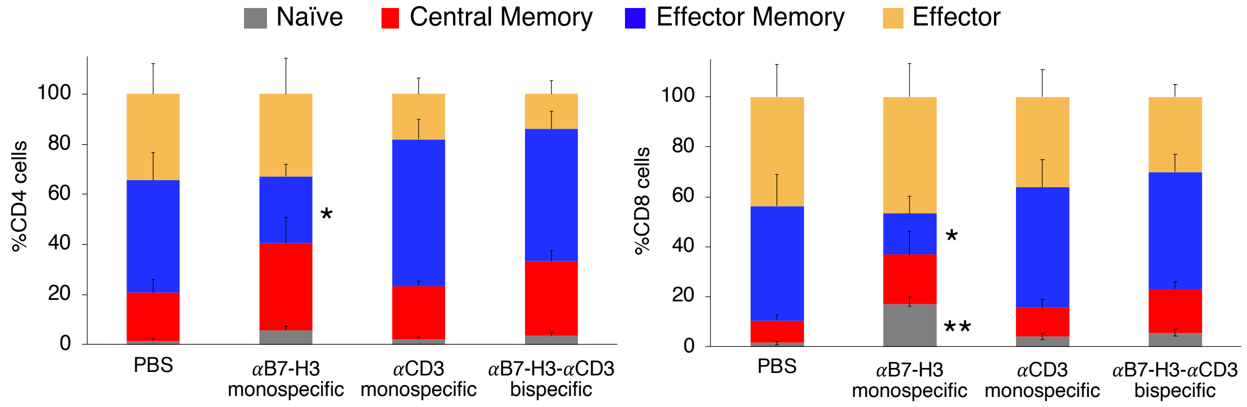


### Supplementary Figure S9.

Systemic memory T cell phenotypes following αB7-H3-αCD3 CSAN treatments *in vivo*. Systemic $CD4^+$ (left) and $CD8^+$ (right) memory cell formation was determined using the lymphocytes collected from peripheral blood and measuring CD45RO and CCR7 expression by flow cytometry. The phenotypes have been previously described as: naïve ($CD45RO^-/CCR7^+$), central memory ($CD45RO^+/CCR7^+$), effector memory ($CD45RO^+/CCR7^-$), effector ($CD45RO^-/CCR7^-$). (*P<0.05 with respect to αB7-H3-αCD3 CSAN treatment against each individual cell phenotype by two-tailed unpaired t-test). Effector and target cells were determined by forward/side scatter and dead cells were removed using a viability dye (n = 5-7 mice per group ± SEM).

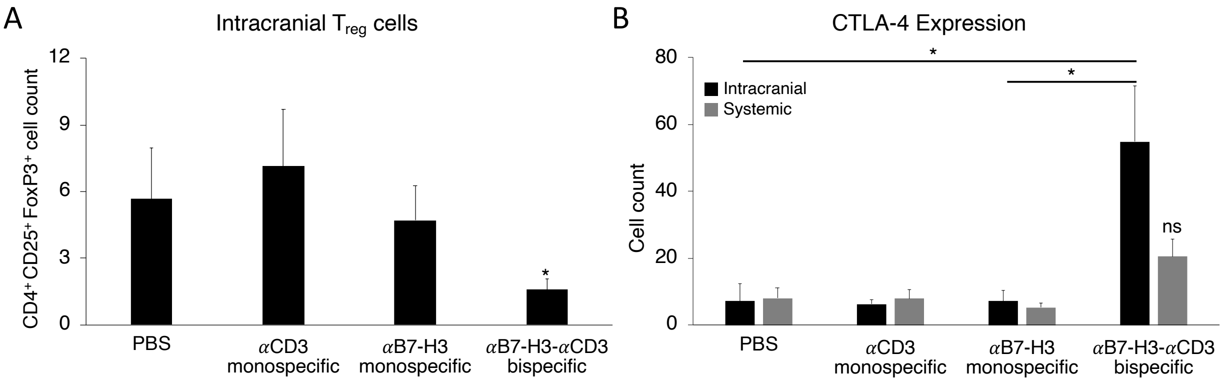


### Supplementary Figure S10.

Bispecific CSAN treatments influence regulatory T cell presence and checkpoint molecule expression *in vivo*. **A,** Presence of Treg cells within the cerebellum single cell suspension was determined by measuring $CD4^{+}/CD25^{+}/FoxP3^{+}$ T cell expression after fixation and permeabilization. **B,** The cerebellum single cell suspension and peripheral blood lymphocytes from each mouse were analyzed for CTLA-4 expression by flow cytometry (Significance is indicated as $^{*}P<0.05$ by two-tailed unpaired t-test). Effector and target cells were determined by forward/side scatter and dead cells were removed using a viability dye (n=5-7 mice per group ± SEM).
